# Pool choice in a vertical landscape: tadpole rearing site flexibility in phytotelm-breeding frogs

**DOI:** 10.1101/2021.03.10.434757

**Authors:** Chloe A. Fouilloux, Shirley Jennifer Serrano-Rojas, Juan David Carvajal-Castro, Janne K. Valkonen, Philippe Gaucher, Marie-Therese Fischer, Andrius Pašukonis, Bibiana Rojas

**Affiliations:** Department of Biology and Environmental Sciences, University of Jyväskylä, PO Box 35, FI 40001, Finland; Department of Biology, Stanford University, 371 Jane Stanford Way, Stanford, CA 94305, USA; Instituto de Investigación de Recursos Biológicos Alexander von Humboldt, Bogotá DC, Colombia; Department of Biological Sciences, St. John’s University, Queens, NY, USA; USR LEEISA - Laboratoire Ecologie, Evolution, Interactions des Systèmes amazoniens, CNRS-Guyane, Cayenne, French Guiana; Centre d’Ecologie Fonctionelle et Evolutive, CNRS, 1919 Route de Mende, 34293 Montpellier Cedex 5, France

**Author notes:** Equal senior author contribution (Andrius Pašukonis, Bibiana Rojas).

**Keywords:** parental care, poison frogs, phytotelmata, niche partitioning, competition, tadpoles

## Abstract

Many species of Neotropical frogs have evolved to deposit their tadpoles in small water bodies inside plant structures called phytotelmata. These pools are small enough to exclude large predators but have limited nutrients and high desiccation risk. Here, we explore phytotelm use by three common Neotropical species: *Osteocephalus oophagus*, an arboreal frog that periodically feeds eggs to its tadpoles; *Dendrobates tinctorius*, a tadpole-transporting poison frog with cannibalistic tadpoles; and *Allobates femoralis*, a terrestrial tadpole-transporting poison frog with omnivorous tadpoles. We found that *D. tinctorius* occupies pools across the chemical and vertical gradient, whereas *A. femoralis* and *O. oophagus* appear to have narrower niches that are restricted primarily by pool height, water capacity, alkalinity, and salinity. *Dendrobates tinctorius* tadpoles are particularly flexible, and can survive in a wide range of chemical, physical, and biological conditions, whereas *O. oophagus* seems to prefer small, clear pools and *A. femoralis* occupies medium-sized pools with abundant leaf litter and low salinity. Together, these results show the possible niche partitioning of phytotelmata among frogs, and provide insight into stressors and resilience of phytotelm breeders.

## Introduction

The survival of young often hinges on the quality of the rearing environments created or chosen by their parents. Whether it is by building nests (birds: (Brown and Brown 1991); mice: (Bult and Lynch 1997, Zhao et al. 2016), digging burrows (rodents: (Svendsen 1976, Ebensperger et al. 2014), or depositing clutches/larvae (e.g., salamanders: (Ruano-Fajardo et al. 2014), frogs: (Pettitt et al. 2018)), the ecology of rearing sites is fundamental in shaping offspring success. For animals with external fertilization, breeding-site choice can be especially important, as optimal conditions for egg clutches may differ from the optimal environment for hatchlings and adults (fish: (Ottesen and Bolla 1998, Mikheev et al. 2001); salamanders: (Nussbaum 1987, Sih and Moore 1993); frogs: (Vági et al. 2019)). Many of these animals assess and prefer biotic and abiotic properties of breeding sites that can enhance offspring survival (Marsh and Borrell 2001, Mokany and Shine 2003, Brown and Shine 2005, Touchon and Worley 2015). Thus, characterizing the nurseries where offspring occur can provide information on the qualities parents assess when making these critical reproductive decisions.

The challenge of finding an optimal rearing site becomes especially apparent in terrestrial or arboreal breeding animals, whose larval forms are aquatic. For example, some treefrogs lay clutches overhanging water bodies. The placement of clutches is essential, as tadpoles from poorly placed clutches risk hatching and falling onto the ground (Wells 2007, Warkentin 2011). One remarkable amphibian strategy adapted to changing habitats between egg and larval stages involves parents that physically transport recently hatched tadpoles from terrestrial oviposition sites to small water-holding plant structures (phytotelmata), ponds, or streams (Summers and Tumulty 2014, Schulte et al. 2020). Unlike other terrestrial breeding amphibians, the physical transport of young allows parents to select the ideal environment for their offspring to develop. Although it is difficult to extensively characterize streams and ponds, microhabitats like phytotelmata provide a unique opportunity to fully measure the biological, chemical, and physical aspects of a nursery, creating an opportunity to interpret deposition choices with a depth of ecological information that is rarely available for other rearing sites. Here, we investigate the chemical and physical properties of aquatic nurseries that predict the presence of Neotropical tadpoles in phytotelm-breeding frogs.

The use of phytotelmata as tadpole nurseries can seem counterintuitive, as their small volume makes them prone to desiccation and limited in food (Summers and McKeon 2004, Summers and Tumulty 2014). However, their small size provides protection from large predators, and various species have evolved different strategies for their offspring to succeed in these pools (substrate specialisation: (von May et al. 2009, Pettitt et al. 2018); trophic egg-feeding: (Weygoldt 1980, Brown et al. 2010); larval aggression/cannibalism: (Poelman and Dicke 2007, Gray et al. 2009, Rojas 2014); pool choice based on specific physical or chemical cues: (Lin et al. 2008, Schulte et al. 2011). Despite the widespread use of phytotelmata, and the non-random site selection shown by many frog parents, few studies go beyond quantifying basic pool dimensions and pool occupation to understand tadpole deposition decisions. Further, the bulk of phytotelm studies are focused only on bromeliads, while work exploring potential trade-offs associated with different phytotelmata (i.e., physical and chemical properties as well as food- and predator-related pressures), and how these change across a vertical gradient, has gone largely overlooked.

To understand what variables drive phytotelm selection, we compared pool occupation by three Neotropical frogs (*Dendrobates tinctorius, Allobates femoralis*, and *Osteocephalus oophagus*) that were most frequently detected in phytotelmata throughout field surveys at our study site in French Guiana. *Osteocephalus oophagus* is a hylid treefrog with bi-parental care and obligately oophagous tadpoles (Jungfer and Weygoldt 1999, Almendáriz et al. 2000). As in our field site, adults have been found to call and breed in bromeliads, tree-holes, and palm axils close to the forest floor (Jungfer and Weygoldt 1999). Tadpoles of this species develop in the same pool in which the eggs are deposited. *Allobates femoralis* is a terrestrial frog closely related to dendrobatids. Adult males aggressively defend territories during the rainy season (Roithmair 1992, Narins et al. 2003), from which they carry recently hatched tadpoles to a variety of terrestrial pools including phytotelmata close to the ground (Ringler et al. 2009, 2013). Tadpoles of this species are omnivorous (McKeon and Summers 2013), but not cannibalistic (Summers and McKeon 2004).

Following broad species-wide comparisons, we focus on a more detailed analysis of pool choice in *D. tinctorius*, a phytotelm specialist with predatory and cannibalistic tadpoles which are deposited in a range of phytotelm types (e.g., palm bracts, tree-holes, fallen trees; Fig. 1 and Fig. 2) that occur from the forest floor to more than 20 m in vertical height (Gaucher 2002, Rojas 2014, 2015). The use of the high canopy pools is perplexing because *D. tinctorius* is commonly successful in terrestrial pools (Rojas 2014). It is known that pool chemistry can change drastically depending on substrate (‘dead’ or ‘live’; see Fig. 1), height, and community composition (Ruano-Fajardo et al. 2014, Pettitt et al. 2018). Thus, better understanding the ecology of high arboreal pools and characterizing phytotelmata across the vertical gradient could help explain both the apparent success of *D. tinctorius* in a wide range of pools, and why parents sometimes carry their offspring such heights. Further, *D. tinctoriu* tadpoles predate on other species of phytotelm-breeders, making it a key amphibian species for understanding the niche partitioning amongst communities of phytotelm breeders. To our knowledge, this is the first study providing detailed biotic, physical, and chemical comparisons of phytotelm choice between Neotropical species.

**Figure 1.**
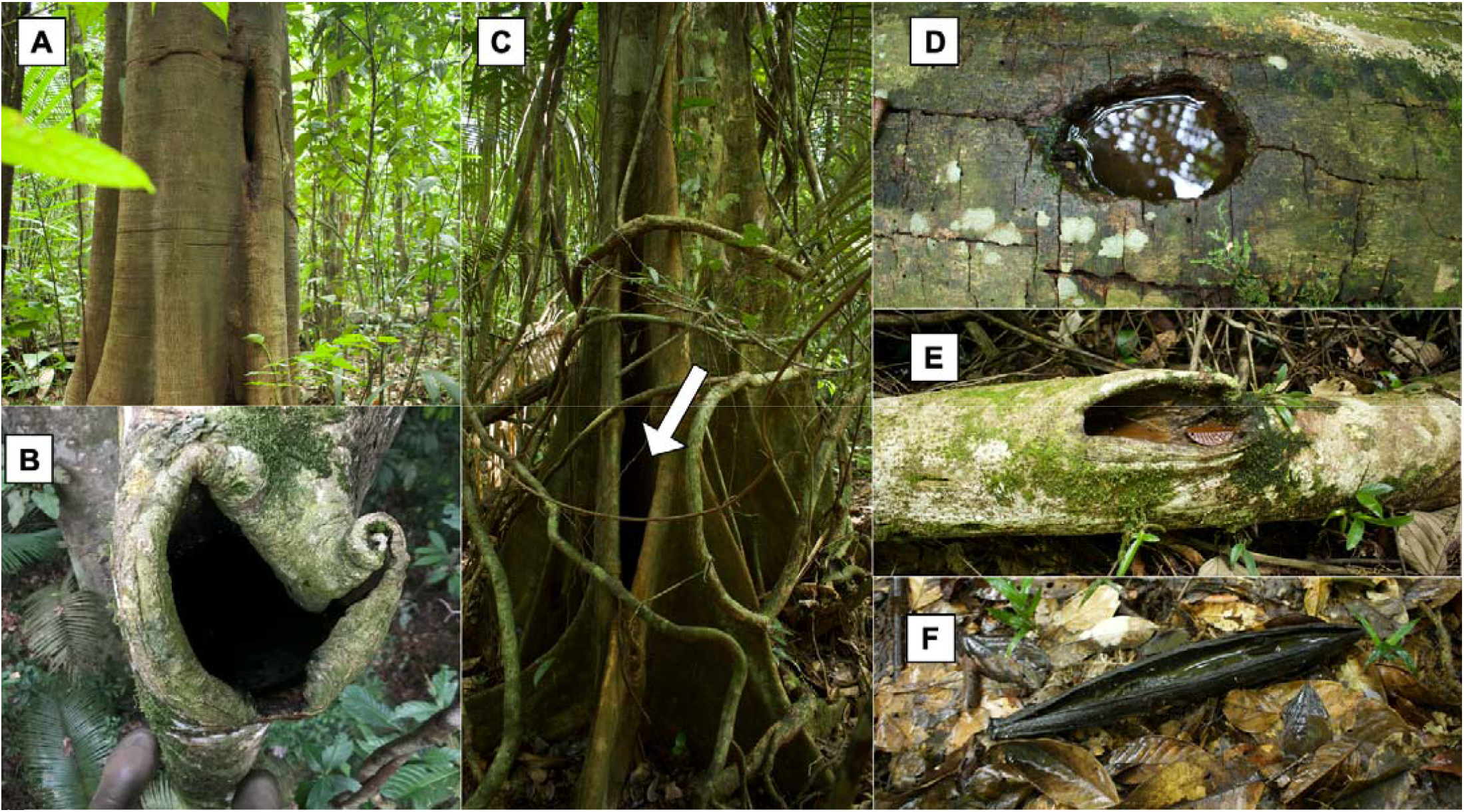
Visual overview of sampled pool diversity. Photos show the diversity of pools across the vertical gradient. Phytotelmata used by frogs include “live” substrate pools such as tree holes **(A)**, high arboreal pools **(B)**, and buttresses **(C)**. There were also commonly occupied “dead” substrate pools such as fallen trees **(D, E)** and palm bracts **(F)**.

**Figure 2.**
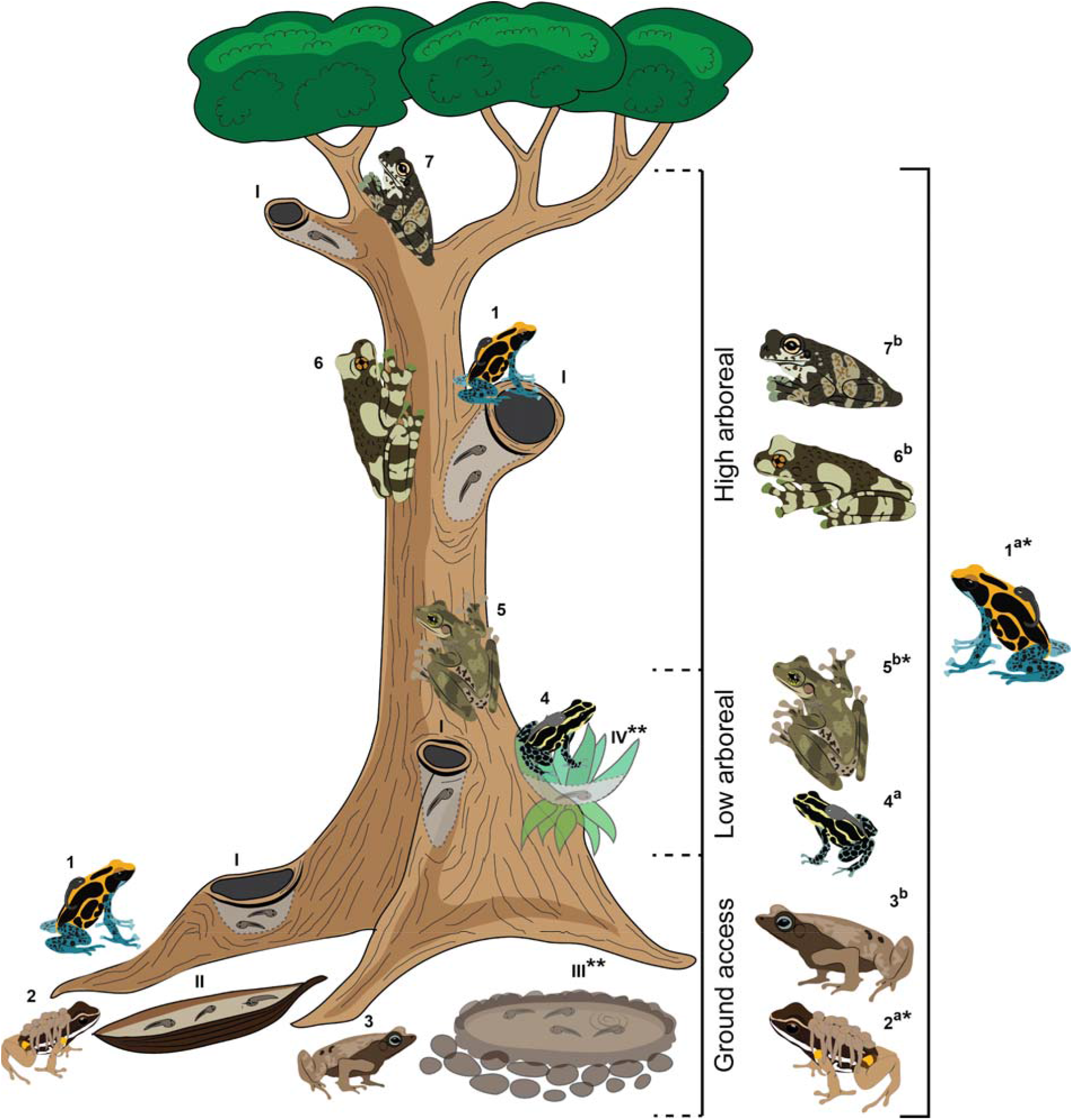
Vertical partitioning of the phytotelm-breeding anuran community in the Nouragues Nature Reserve, French Guiana. Numbers indicate seven species detected during this study: 1. Dendrobates tinctorius, 2. Allobates femoralis 3. Rhinella castaneotica, 4. Ranitomeya amazonica, 5. Osteocephalus oophagus, 6. Trachycephalus resinifictrix, 7. T. hadroceps. Letters indicate species with a) terrestrial or b) aquatic eggs. Latin numerals indicate commonly used pool types: I. tree holes at various heights, II. fallen palm bracts, III. ground puddles, IV. bromeliads. *Most commonly encountered species included in the analysis; ** Pool types not sampled in this study (see methods).

## Material and methods

The study was carried out in the primary lowland terra-firme forest near the Camp Pararé at the CNRS Nouragues Ecological Research Station in the Nature Reserve Les Nouragues, French Guiana (4°02’N, 52°41’W) over two field seasons: 1st February to 20th March 2019, and 30th January to 26th February 2020. The study area was chosen specifically because of the high abundance of *D. tinctorius* (*Rojas and Pašukonis 2019*). Pools were found with a combination of field methods. We opportunistically searched for pools targeting suitable microhabitats such as fallen trees, trees with buttresses and palm trees. In addition, pools were discovered by using tracking to follow *D. tinctorius* during previous studies (Pašukonis et al. 2019). We also used experimentally-induced tadpole transport in combination with tracking (Pašukonis et al. 2017) to find additional pools used by *A. femoralis*. Trees with high arboreal pools were discovered by locating calls produced by the treehole-breeding frogs *Trachycephalus resinifictrix* and *T. hadroceps* during night surveys.

### Sampled pools

We exclusively considered phytotelmata throughout this study. Pools could be classified into two categories: dead substrates, which included holes in dead branches, fallen trees, and fallen *Oenocarpus* palm bracts; or live substrates which included live tree trunks, branches, roots, and buttresses. We did not sample bromeliads and non-phytotelm pools as these pools are not used by *D. tinctorius*. Based on the pools’ height and accessibility to different frog species, we termed the pools as “ground access”, “low arboreal” or “high arboreal” (Fig. 1, 2). Ground-access pools did not require vertical climbing ability to reach and included dead fallen structures as well as pools in live roots or low buttresses. Low arboreal pools were inside vertical structures low on the trunk or on high buttresses. High arboreal pools were high on the trunk or in canopy branches and were accessed for sampling using rope-based canopy access methods. There was a clear vertical separation between ground-access and low-arboreal pools, which were all under 212 cm in height and between those and high arboreal pools, which were all above seven meters in height. In total, we sampled 84 unique pools across the 2019 and 2020 field seasons.

Several unique pools were sometimes found and sampled in the same tree. For all pools, we recorded the pool type, location (latitude/longitude), height from the ground to the pool edge, largest width and length parallel to the water surface, and the pool depth from the solid sediment bottom to the maximum water level line. Based on these measurements, we estimated the maximum water-holding capacity of each pool using the volume formula of a semi-ellipsoid as in Rojas (2014). Other sampling methods differed between the two field seasons.

### 2019 field season sampling

In 2019, we quantified physical measures (height, pool dimensions, leaf-litter volume), biotic measures (amphibian and invertebrate counts and diversity), as well as chemical measurements (see Supp. Table 1 for description of all variables measured). For pools accessible from the ground and smaller arboreal pools, we attempted to sample all tadpoles and Odonata larvae (primary tadpole predators) in each pool. Initially, we carefully observed the undisturbed pool and attempted to catch all tadpoles and Odonata larvae using a variety of tools. We then syphoned the entire volume of the water and sediment from the pool, emptied the leaf litter and searched for tadpoles and Odonata larvae. The volumes of water, sediment, and leaf litter were measured. For deep arboreal pools, we repeatedly netted and observed the pool until no more tadpoles were caught during five minutes of continuous netting. We carefully scraped the inner walls of the pools and caught as many Odonata larvae as possible. The leaf-litter volume could not be accurately measured for some deep arboreal pools, but they typically were protected from falling leaves and had little leaf litter in them.

**Table 1.** Principal component regression of tadpole presence in phytotelm pools. Using a negative binomial GLM, we found that only the first principal component is significant in predicting tadpole presence.

| Tadpole presence (Y/N) |  |  |  |
| --- | --- | --- | --- |
| *Predictors* | *Estimates* | *CI* | *p* |
| (Intercept) | -0.85 | -1.27 – -0.49 | **<0.001** |
| PC1 | 0.25 | 0.08 – 0.42 | **0.003** |
| PC2 | 0.09 | -0.16 – 0.40 | 0.528 |
| PC3 | 0.21 | -0.07 – 0.48 | 0.144 |

We used visually apparent morphological traits to identify tadpoles, except for *Allobates femoralis, A. granti, and Ameerega hahneli*, which we could not reliably differentiate in the field. Because *Allobates femoralis* was more common in our study area than *A. granti* and *Am. hahneli* and we never observed *A. granti* and *Am. hahneli* directly at the pools we classified all *A. femoralis-*like tadpoles as such. Is it important to note that some *A. granti* and *Am. hahneli* tadpoles may have been misclassified as *A. femoralis*. However, this does not affect the interpretation of our results as all three species are cryptic terrestrial poison frogs similar in appearance, ecology and behavior. We also opportunistically recorded all species of adult frogs heard or seen at each pool throughout the sampling period. This was used as an amphibian diversity index between 0 and 8 species observed at each pool. Tadpoles of only three out of seven recorded species, namely *D. tinctorius, O. oophagus* and *A. femoralis*, were detected in pools with sufficient frequency for further analysis (N = 34 (2019), N = 7 and N = 10 pools, respectively).

Sampled invertebrates were counted, photographed, and classified only to a group level (usually order or class) apparent in the field. To estimate the predation pressure on tadpoles we used the total count and average size of all Odonata larvae detected in the procedure described above. To estimate density and diversity of aquatic invertebrates, we carefully searched and counted invertebrates in a sample of up to 10 liters of water and up to one liter of sediment in proportion to the total estimated pool volume. For each liter of the water volume sampled, we sampled ∼ 100 mL of sediment from the bottom of the pool. When the water volume was less than one liter or the amount of sediment was less than 100 mL, we sampled the entire pool and recorded the exact volumes. In the final analysis, we used the invertebrate density (count divided by the volume sampled) and the diversity index corresponding to our classification (between 0 and 12). The following 12 categories were used to quantify invertebrate diversity: Odonata Zygoptera larvae, Odonata Anisoptera larvae, surface Coleoptera adults, diving Coleoptera adults, Coleoptera Scirtidae larvae, Trichoptera larvae, Diptera Culicidae larvae, Diptera Chironomidae larvae, Diptera Tipulidae larvae, other Diptera larvae, small red Annelida, other unidentified larvae. All water, sediment, tadpoles and invertebrates were released back into the pool after sampling.

We measured water conductivity, salinity and total dissolved solids (TDS), dissolved oxygen and temperature with electronic sensors (EZDO 7200 and pHenomenal OX4110H). Water chemistry (KH (also known as alkalinity), hardness and NO_3_) was recorded using aquarium water testing strips (JBL EasyTest). All measures were taken from the undisturbed surface water of the pool.

### 2020 field season sampling

The 2020 dataset focused solely on *D. tinctorius* tadpole counts and pH measurements of weekly resampled ground access phytotelmata (N = 26) over the time period of a month (February 2020). Rainfall data were provided by the Nouragues Ecological Research Station from an above-canopy weather station in the study area. High arboreal pools (N = 8 (2020)) were only measured once. pH was recorded using a pH meter (AMTAST Waterproof pH Meter). The pH meter was calibrated once per day, prior to pool sampling, using both acidic (pH = 4) and neutral (pH = 7) calibration solutions. The pH of ground access pools was taken directly by submerging the pH probe into the pool, and the measurement was recorded once read-out stabilized. For arboreal pools, a sample of water was collected using a syringe, which was then sealed at both ends. Once on the ground, one end of the syringe was opened, and the pH was measured by submerging the pH probe into the syringe. Syringes were never reused. Between pool sampling, the pH probe was wiped with a clean cloth and rinsed with aquifer water.

### Statistical Analyses

The presence of *D. tinctorius* in pools was analyzed using 2019 field data. As a result of the high collinearity between variables in the 2019 dataset (see Supp Fig 1), we used a principal components regression to analyze phytotelm ecology data. We first checked data for a non-random structure following Björklund (Björklund 2019); then, we established that the correlation matrices were significantly different from random (**Ψ**= 10.22, *p* = 0; *ϕ*= 0.238, *p* < 0.001) to ensure they were suitable for a PCA. Based on each PCs difference from random matrices, we selected the first three principal components as predictors of probability for *D. tinctorius* tadpole presence as a binomial response in the principal component regression (PC1-3 explained about 53% of the variation proportion of variance explained ± SE : PC1 = 0.24 ± 0.48, PC2 = 0.17 ± 0.40, PC3 = 0.11 ± 0.33). We evaluated the fit of negative binomial GLM models based on AIC ranks ((Akaike 1974); see Supp. Table 2). Models within two AIC scores of each other were further evaluated by assessing the significance of interactions between model terms.

**Table 2.**
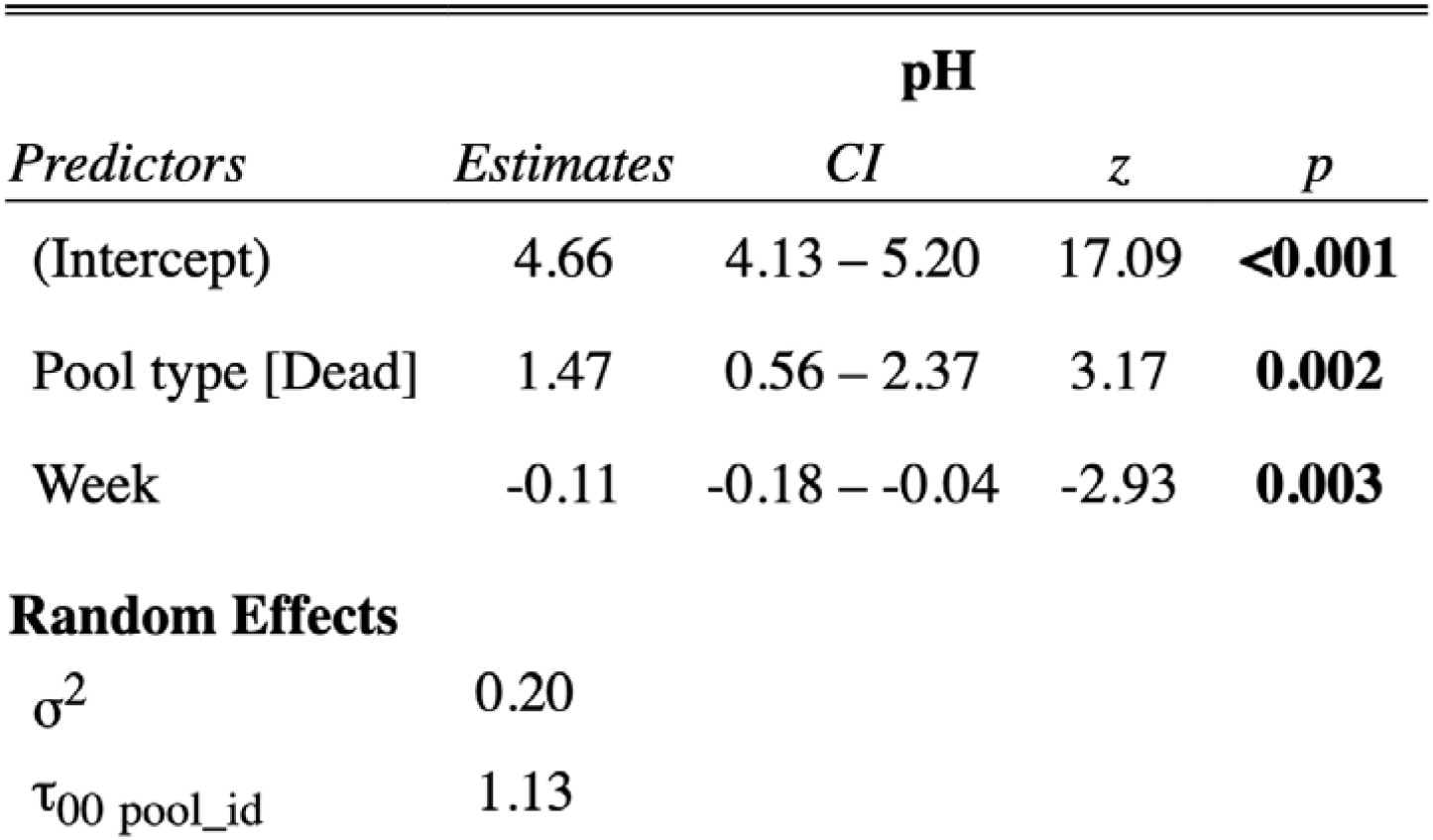
Negative binomial generalized mixed model predicting pH over time based on pool substrate. Data include low arboreal and ground access pools. Pool type is a 2-level categorical variable (“Dead”, “Live”). Where pH is significantly higher in dead pool type versus live pools type, and decreases over time.

| pH |  |  |  |  |
| --- | --- | --- | --- | --- |
| Predictors | Estimates | CI | z | p |
| (Intercept) | 4.66 | 4.13 – 5.20 | 17.09 | **<0.001** |
| Pool type [Dead] | 1.47 | 0.56 – 2.37 | 3.17 | **0.002** |
| Week | -0.11 | -0.18 – -0.04 | -2.93 | **0.003** |
| **Random Effects** |  |  |  |  |
| $\sigma^2$ | 0.20 | | | |
| $\tau_{00}$ pool_id | 1.13 | | | |

To better understand which variables contributed significantly to each principal component, we calculated which variables had index loadings larger than random data. Following the methods outlined by Björklund (2019) and Vieira (2012), we randomized the data and calculated new correlation matrices which we permuted 1,000 times. We then compared the index of loadings (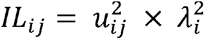, see Vieira (2012) for details) between each observed PC and the randomly generated data to see which variables contributed significantly to each principal component.

The 2020 dataset consisted of weekly resampled pools throughout February 2020. Models took repeated measures of pool ID into account as a random effect. Both the presence of *D. tinctorius* tadpoles (count; negative binomial family) and pH (Gaussian family) from resampled pools were modeled using a mixed effects generalized linear model in the package “glmmTMB” (Magnusson et al. 2020). Predictor structure for both pH and *D. tinctorius* models were built based on biologically relevant variables (pool substrate, time, *D. tinctorius* tadpole count (for pH model), water capacity, surface area:depth ratio). Using these variables, models were first fit with relevant interactions (see Supp. Table 3, 4), which were then removed if they did not contribute significantly to the model using single term deletions (drop1). Residuals were diagnosed using the package “DHARMa” (Hartig 2020); final models were then checked for overdispersion and zero-inflation and corrected as necessary. All code was done in R (Core 2015).

**Table 3.** Negative binomial generalized mixed model predicting D. tinctorius tadpoles in resampled pools in February 2020.

| *D. tinctorius* tadpoles (count) |  |  |  |  |
| --- | --- | --- | --- | --- |
| *Predictors* | *Estimate* | *CI* | *z* | *p* |
| (Intercept) | -10.35 | -16.65 – -4.05 | -3.22 | **0.001** |
| Pool type [Dead] | 11.08 | 2.96 – 19.21 | 2.67 | **0.008** |
| pH | 1.59 | 0.48 – 2.71 | 2.80 | **0.005** |
| Week | -0.16 | -0.39 – 0.06 | -1.40 | 0.160 |
| Pool type [Dead]: pH | -1.50 | -2.93 – -0.07 | -2.05 | **0.040** |
| **Random Effects** |  |  |  |  |
| $\sigma^2$ | 2.62 | | | |
| $\tau_{00}$ pool_id | 1.87 | | | |

## Results

### Species-wide trends

We detected tadpoles or adults of 7 species from 4 families using surveyed phytotelmata for breeding (Fig. 2). Dendrobatidae: *Dendrobates tinctorius, Ranitomeya amazonica*, Aromobatidae: *Allobates femoralis*; Hylidae: *Osteocephalus oophagus, Trachycephalus resinifictrix, T. hardroceps*; Bufonidae: *Rhinella castaneotica*. The tadpoles of only three species (*D. tinctorius, O. oophagus* and *A. femoralis*, present in N = 34, N = 7 and N= 10 pools, respectively) were detected frequently enough for further analysis. Although the data on *D. tinctorius* are more robust, trends for *O. oophagus* (N = 125 tadpoles) and *A. femoralis* (N = 117 tadpoles) emerge despite a smaller dataset. The species-wide dataset is based on the sampling of 70 unique pools in 2019.

Differences in pool accessibility are highlighted by Fig. 3. Compared to *A. femoralis* and *O. oophagus*, one of the most striking aspects of *D. tinctorius* ecology is its flexibility with respect to site choice on a vertical axis. *Dendrobates tinctorius* tadpoles were found in pools from the forest floor to more than 15 meters in the canopy. For *O. oophagus*, a strictly arboreal frog in its adult stage, tadpoles were detected only in low arboreal pools, ranging from 20 cm to 1.7 m in height. In *A. femoralis*, tadpoles were only found in ground-access pools, and occurred at a maximum height of 71 cm.

**Figure 3.**
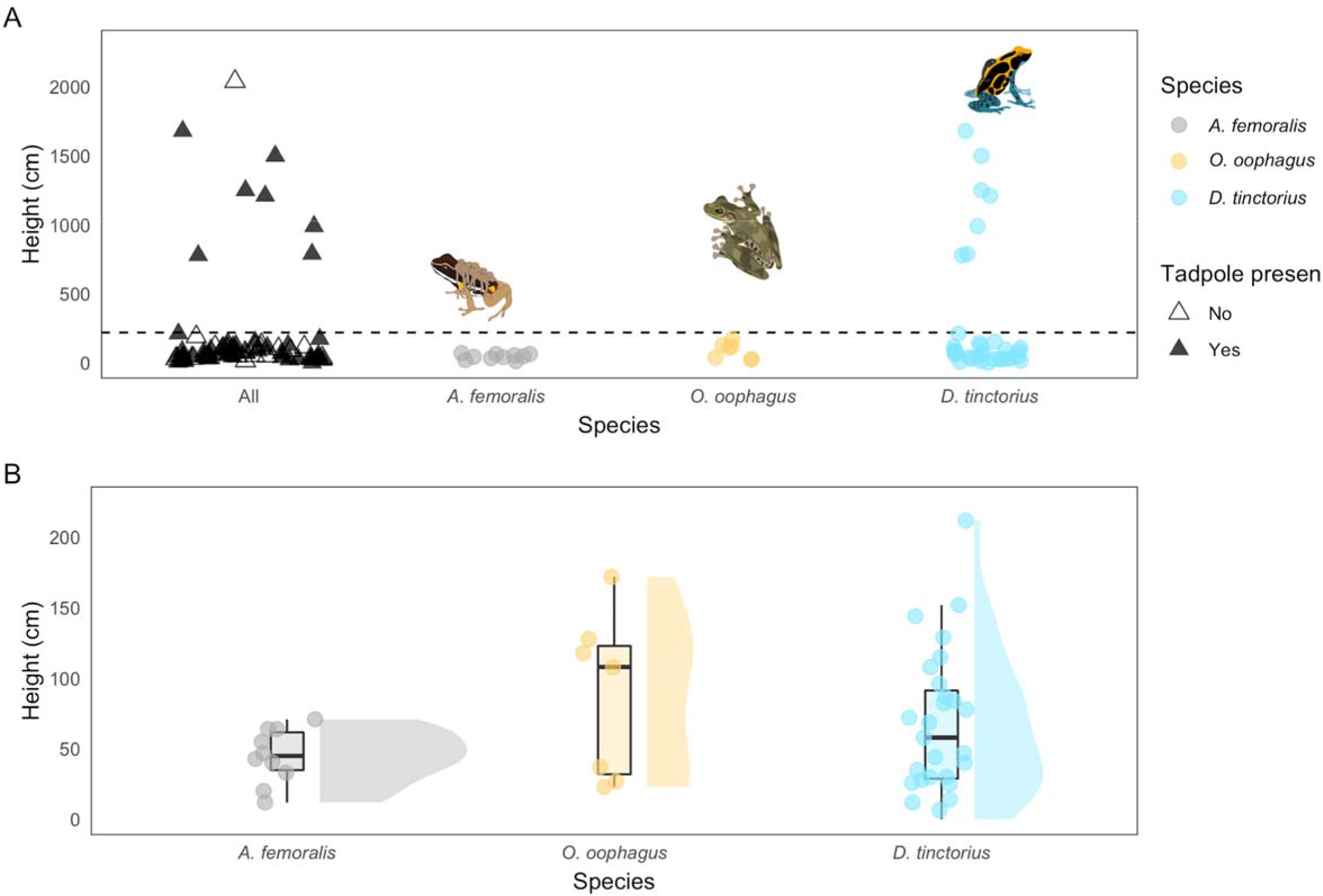
Tadpole presence across the vertical landscape. Panel **(A)** shows all sampled pools. For the “All” category, colored/empty triangles represent presence/absence data of all three species. Dashed line is drawn at 220cm; pools above this limit are classified as high arboreal pools. Panel **(B)** highlights occupied pools below 220 cm. Dendrobates tinctorius (N= 34) tadpoles occur in pools across the vertical landscape. Distribution of O. oophagus (N = 7) and A. femoralis (N = 10) tadpoles shows possible vertical niche partitioning. Boxplot whiskers extend 1.5 * interquartile range. Violin plots represent density distribution for species occurrence. Data is from the 2019 field season.

Despite small sample sizes we found clear trends: *O. oophagus* tadpoles are heavily biased towards small, clear pools and *A. femoralis* is present in medium and large pools with large amounts of leaf litter, whereas *D. tinctorius* occurs throughout the sampled range (Fig. 4). As opposed to *A. femoralis* and *O. oophagus, D. tinctorius* can occupy chemically diverse pools, showing remarkable flexibility with respect to KH, salinity, and hardness that clearly limits the other species. *Allobates femoralis* and *O. oophagus* appear to exist in similar KH ranges (KH = 3-6), while *D. tinctorius* appears more tolerant of extreme values (KH = 3-20). *Allobates femoralis* tadpoles occurred in pools with a salinity range from 5 to 37 ppm, while *O. oophagus* tadpoles occupied pools with a range from 48 to 225 ppm (Fig. 5, Panel C). *Dendrobates tinctorius* again appears to have no functional limitation, occupying pools with salinity from 11 ppm up to 955 ppm.

**Figure 4.**
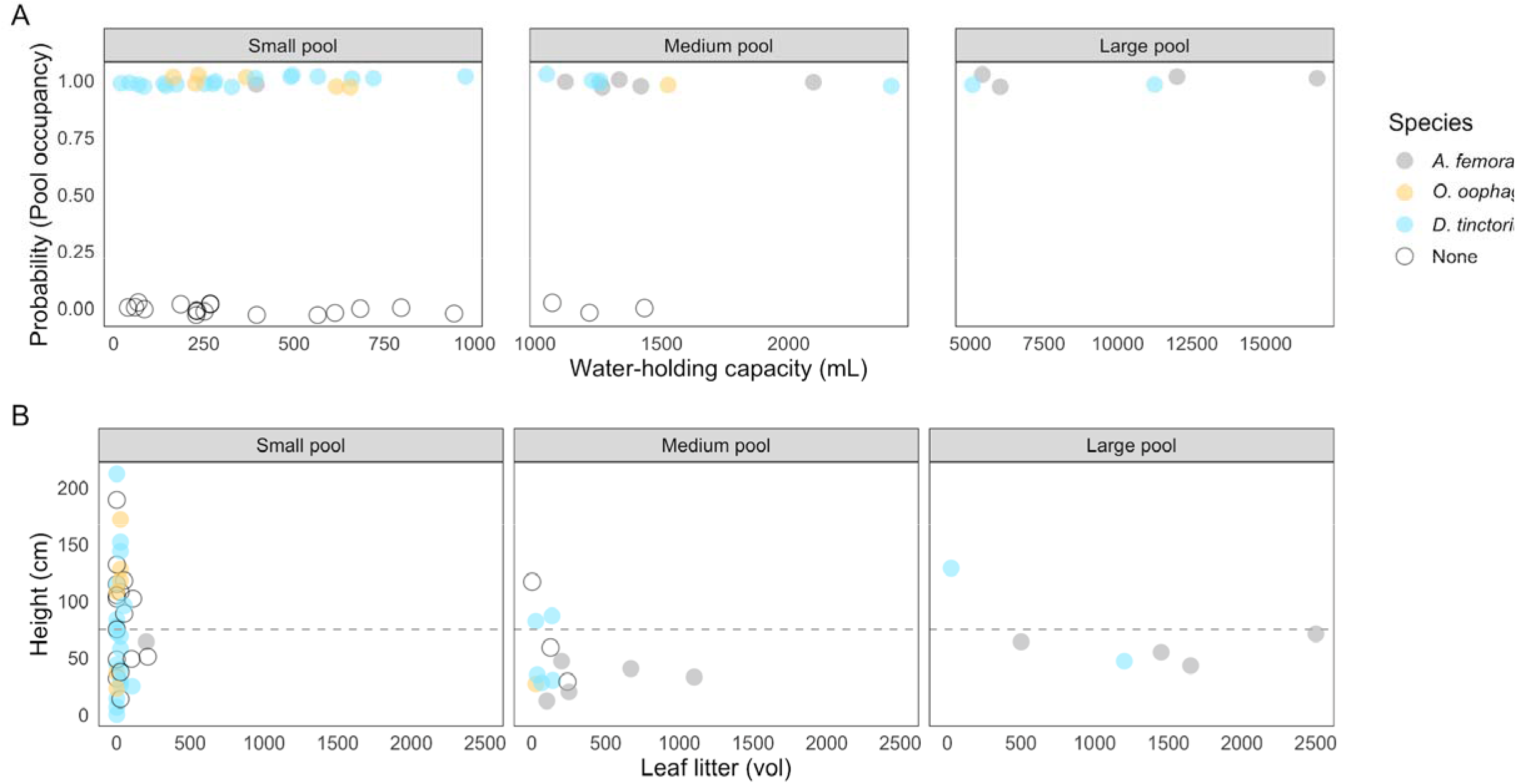
Pool occupancy based on water capacity and leaf-litter volume of phytotelmata. All data are subsetted for low arboreal and ground-access pools (< 220 cm). Panel (**A**) is the probability of pool occupancy (binomial, 0/1) based on water capacity; data are faceted based on relative pool size (Small = < 1000 mL, Medium = < 5000 mL, and Large = > 5000 mL). Points are plotted with a small amount of random noise on the y-axis to facilitate visualization of overlapping data. Panel (**B**) illustrates the correlation between leaf litter and height, faceted by the same pool categories as Panel A. Points are colored by species presence. Dashed line indicates the vertical limit of A. femoralis (< 75 cm). Out of the 62 ground-access and low arboreal pools observed, D. tinctorius co-occurred once with A. femoralis and once with O. oophagus; O. oophagus and A. femoralis tadpoles were never found in the same pool.

**Figure 5.**
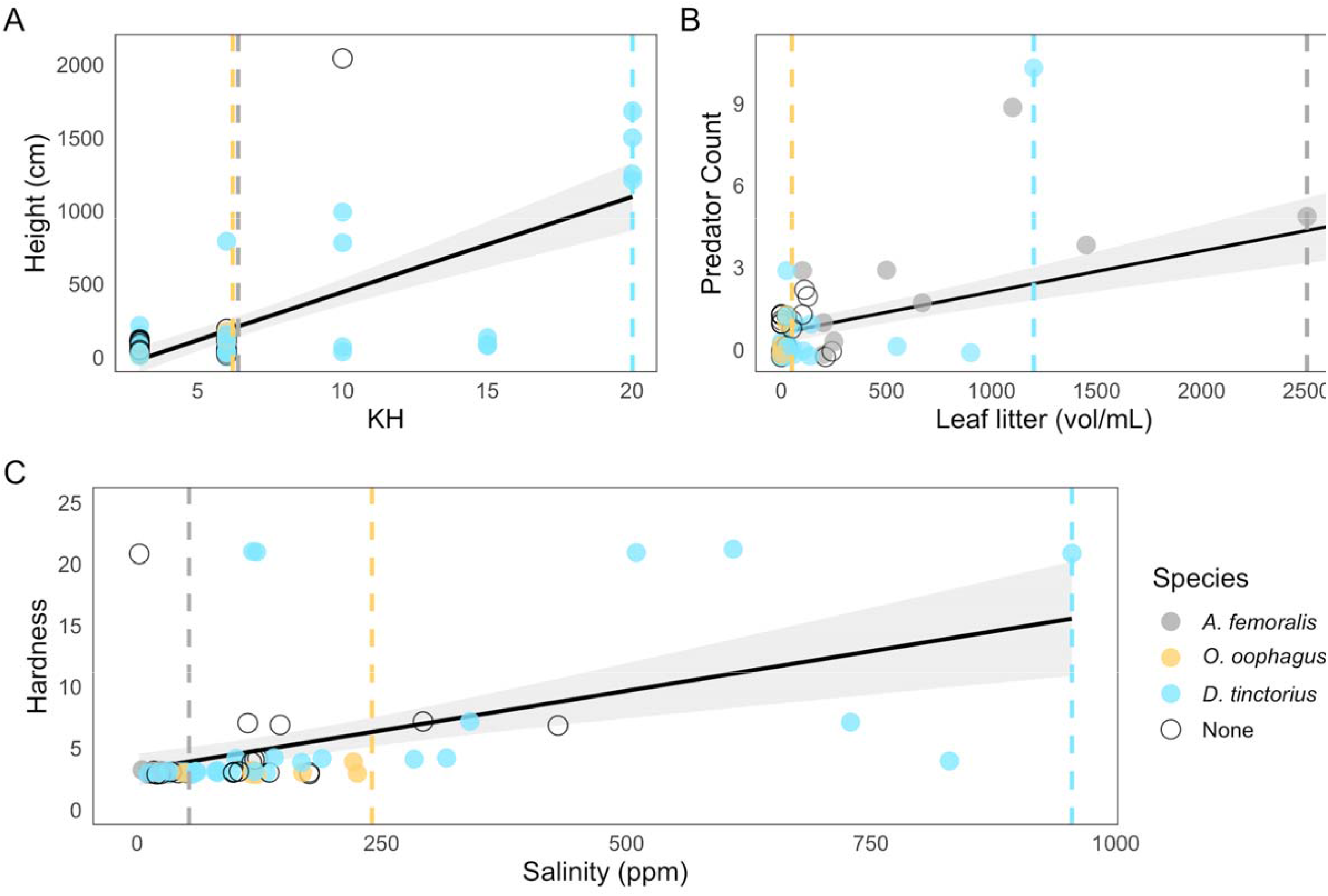
Chemical and physical predictors of tadpole presence in Neotropical tadpoles. We find that D. tinctorius tadpoles are tolerant to a wide range of KH, hardness, height, and salinity values, but appear to be limited with respect to high leaf litter volumes. Plots are based on variables with varied species limits (dashed lines). Colored points represent species presence. Black lines are fit with a GLM smoother, with 95% CI highlighted in light grey.

### Deposition site decisions: *Dendrobates tinctorius*

Because we detected *D. tinctorius* tadpoles much more frequently, we were able to conduct a more thorough analysis of the variables predicting tadpole presence in this species (see Supp. Table 1). We used principal components as predictors for *D. tinctorius* presence. Based on an AIC model comparison, we did not detect any significant interactions between components (Supp. Table 2). A negative binomial GLM only detected PC1 to play a significant role in predicting tadpole presence (Table 1, CI: 0.08-0.42, p = 0.003), where an increase in component value increased the probability of detecting tadpoles.

Following Björklund (2019), we found that, when compared to randomly generated matrices, five out of the original 14 traits (see Supp. Table 1 for trait definitions) contributed significantly to the first principal component. The significant traits can be broadly categorized using three descriptors: (1) chemical (KH, p < 0.001; IL = 1.50, hardness, p = 0.001, IL = 1.30; salinity, p < 0.001, IL = 1.62); (2) physical (height, p = 0.013, IL = 1.06); and (3) biological (invertebrate diversity, p < 0.001, IL = 1.20) (see Fig. 6). Altogether, these results show that *D. tinctorius* tadpoles were found significantly more frequently in pools with higher levels of hardness, KH, and salinity; higher in the vertical gradient; and with more diverse invertebrate communities (Fig. 6).

**Figure 6.**
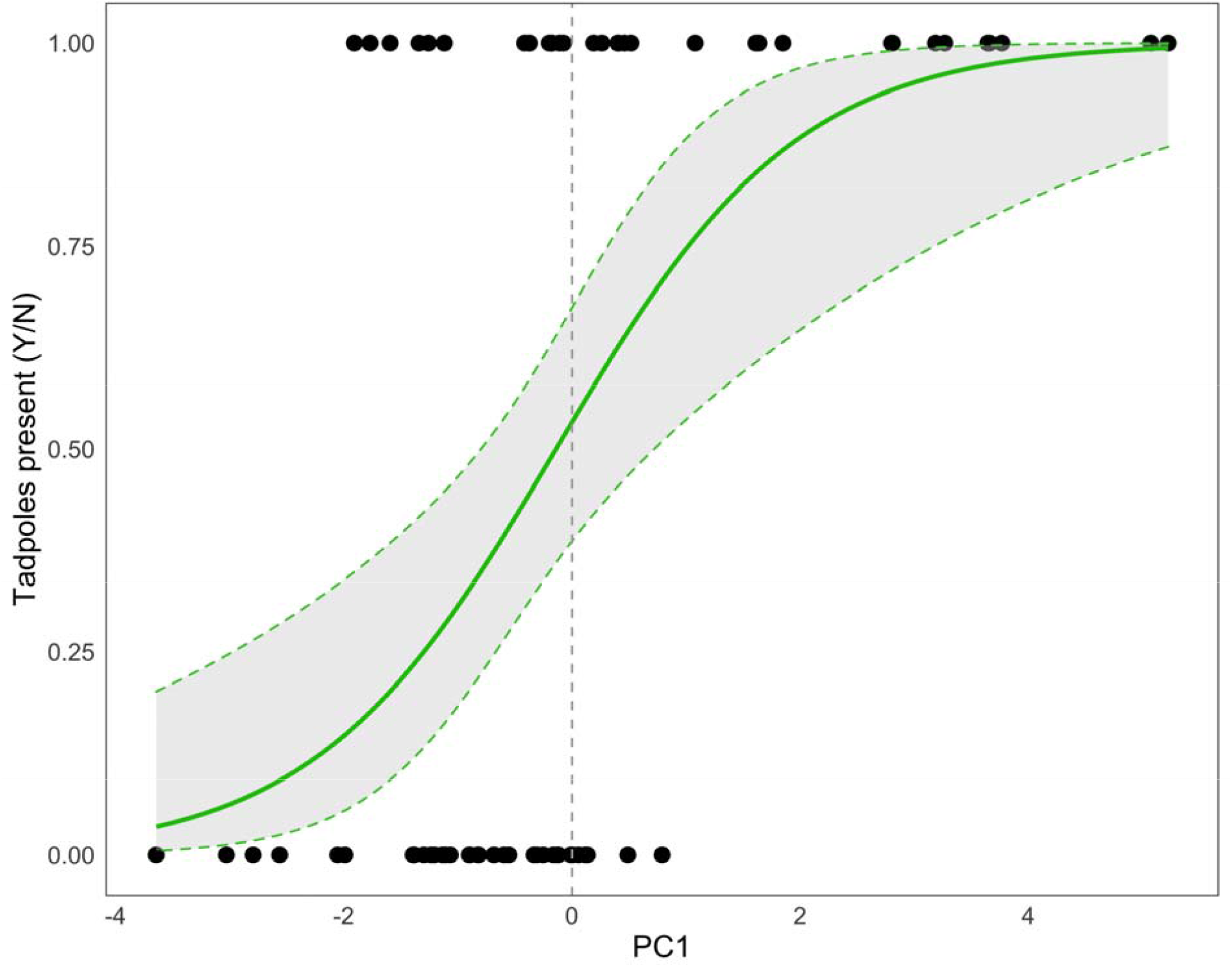
Binomial response of Dendrobates tinctorius presence to the first principal component. Dashed lines represent the 95% confidence interval. Significant variables within this component are detailed in Supp. Table 1.

### *Dendrobates tinctorius* across temporal scales

Using both 2019 and 2020 datasets, we were able to follow phytotelmata across multiple timescales: 13 weekly resampled ground-access and low arboreal pools, 13 annually resampled ground-access and low arboreal pools, and 7 annually resampled high arboreal pools. Overall, we found that pools can persist over multiple sampling seasons. High arboreal pools appear to be the most stable with respect to both tadpole count and tadpole density compared to low arboreal and ground-access pools sampled both years (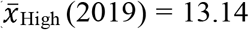 tadpoles, 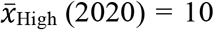 tadpoles versus 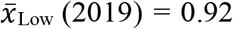 tadpoles, 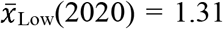). High arboreal pools also had the highest average pH and KH (pH_High_ = 6.73, KH_High_ = 15.14) compared to averages of other pool substrates (pH_(Low)Live_= 4.35, pH_(Low)Dead_= 5.68; KH_(Low)Live_ = 5.69, KH_(Low)Dead_ = 5.88). Due to difficult accessibility, high arboreal pools were sampled only once per year, and thus were excluded from further analysis involving repeated sampling.

When considering pools resampled weekly over the course of a month, two trends emerge: (1) pH is consistently higher in pools contained in ‘dead’ substrates than in ‘live’ substrates (*Odds Ratio*= *1*.*47, Table 2*). For all substrate types, however, pH values decreased over the one-month sampling period (Fig. 7A; *Odds Ratio* = *-0*.*11, Table 2*, which may be related to rainfall levels throughout the month); and (2) the number of *D. tinctorius* tadpoles can be predicted, in part, by the interaction between pool substrate and pH (Fig.7B). Dead pools have higher numbers of *D. tinctorius* tadpoles (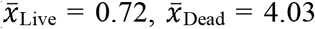, 2020 data). This pH/substrate relationship is clearly important, as tadpoles occur in higher numbers in high pH pools. Time (in weeks) was not detected as an important variable in determining *D. tinctorius* tadpole numbers throughout the month.

**Figure 7.**
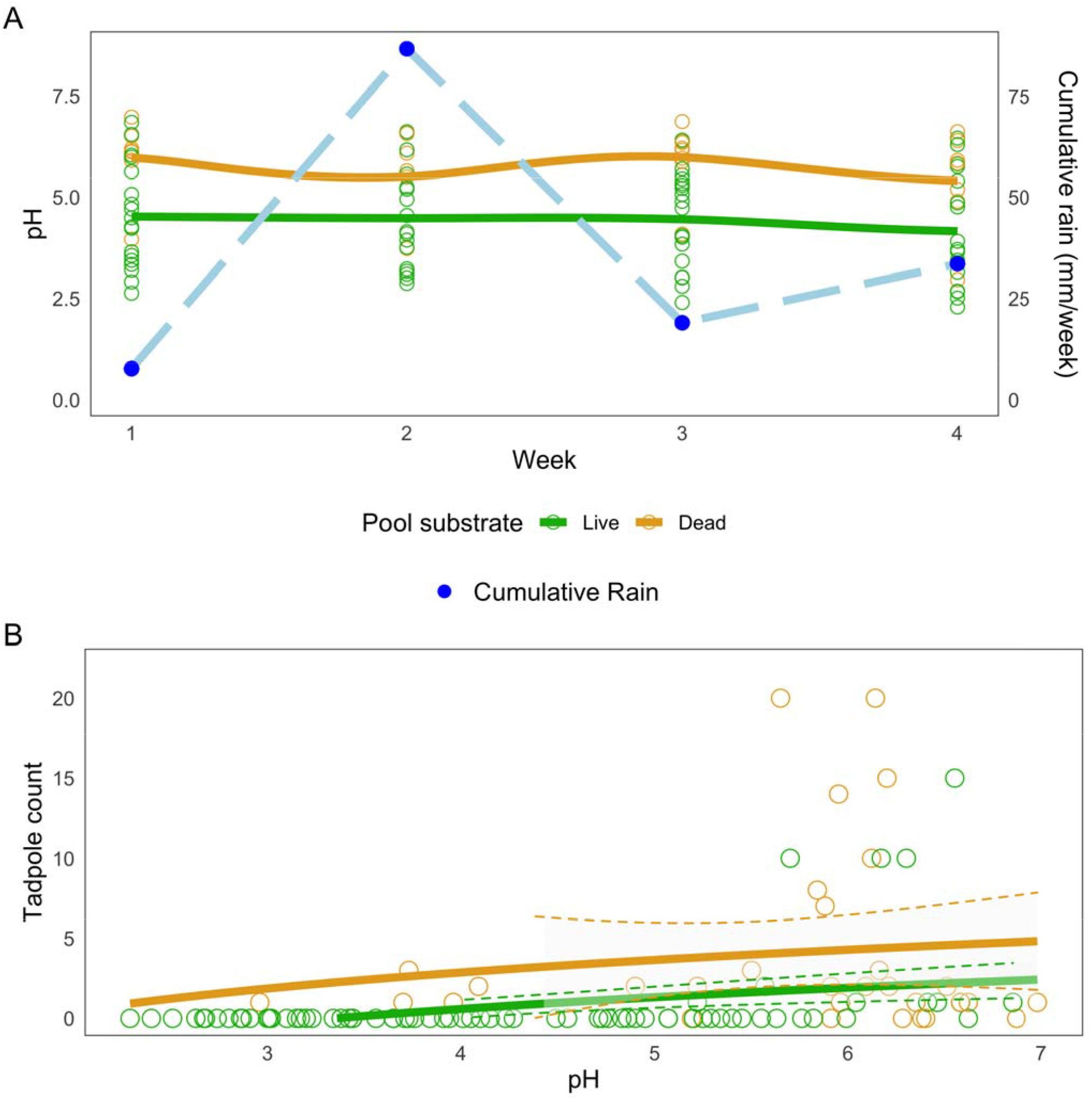
Phytotelm stability and tadpole presence across a month. Panels **(A, B)** are subsetted to exclude high arboreal pools, and emphasize how pH is related to **(A)** pool substrate and **(B)** predicting D. tinctorius tadpole presence in resampled pools. Phytotelmata made of dead substrate have higher pH values than live substrate (N_Dead_ = 8, N_Live_ = 18;each resampled four times) (Table 2); more tadpoles are found in pools with a higher pH (Table 3). Dashed lines represent 95% CI. Blue points in **(A)** indicate the weekly accumulation of rainfall (February 2020). Tan lines are dead substrates and light green line are live substrates; lines in **A** are plotted with a “LOESS” fit and **B** with a “GLM” fit.

Several pools (N = 5, 2020) dried out completely during the month-long observation period, three of which were fallen palm bracts. Thus, although dead phytotelmata tend to have higher pH values, and have a remarkable buffering capacity when filled, they also appear to dry out more easily during dry periods.

## Discussion

Juvenile stages of development are particularly deadly for animals with external fertilization. Thus, the decision of where to breed and raise young is vital to offspring survival. In this study, we investigated the tadpole rearing-site ecology of Neotropical phytotelm-breeding frogs. Out of seven detected species, five showed some form of parental care, three out of which were sufficiently common to infer patterns of pool choice. Broadly, we found that the deposition choices of two of the three species are characterized by height and pool size, such that *O. oophagus* tadpoles occur in small, low arboreal pools below ∼ 2 m and *A. femoralis* tadpoles occur in medium and large pools below ∼ 1 m and with access only from the ground, which is not surprising considering that adults are poor climbers (Roithmair 1992). *Dendrobates tinctorius* tadpoles, in contrast, occur in pools from the ground to the canopy, and of sizes ranging from 19.6 mL to 270L. When focusing on *D. tinctorius* pool choice, we found that despite being able to tolerate an impressive range of physical/chemical factors, tadpoles are more likely to be found in higher pools with greater levels of KH, salinity, and hardness, and higher invertebrate diversity.

### Inter-specific comparison of rearing-site choices

In line with previous research, we found that the preference for water capacity varies among species, and that some of this variation is explained by parental behavior (Lin et al. 2008, McKeon and Summers 2013, Summers and Tumulty 2014). For example, *O. oophagus* provisions their tadpoles with trophic eggs (Jungfer and Weygoldt 1999), which allows parents to choose very small arboreal phytotelmata with rather clear water and little food (Brown et al. 2008a, b, 2010, Summers and Tumulty 2014). Despite the desiccation risk associated with the notably small pools chosen by *O. oophagus*, their nurseries are predator-free, which is often suggested as the primary factor driving the invasion of phytotelmata (Crump 1974, Magnusson and Hero 1991, Gomez-Mestre et al. 2012, Summers and Tumulty 2014).

In contrast, *Allobates femoralis* does not provision tadpoles and preferably transports them to medium-to-large ground access pools. These pools tend to have both high leaf-litter concentrations and a high number of predators (Fig. 5B; concurrent with McKeon and Summers (2013)). The effect of leaf litter on Neotropical tadpoles is unclear, but large amounts have been found to have both positive effects (increased growth rate in African tadpoles; (Lehtinen 2004)) and negative effects (decreased growth rate and development in temperate-region tadpoles; (Williams et al. 2008)). Because *A. femoralis* are confined to ground access phytotelmata due to their inability to climb, choosing to deposit their tadpoles in pools containing high amounts of leaf litter may be their best option: despite the higher predation risk (which *A. femoralis* fathers try to minimize (Ringler et al. 2018)), leaf litter provides a source of food and shelter/predator protection to tadpoles that do not exist in clear pools. Interestingly, the turbid leaf litter pools occupied by *A. femoralis* were functionally available to *D. tinctorius*, who do not appear to use them. As *D. tinctorius* tadpoles are predatory, clear pools may be better suited for visual foraging.

Contrary to our expectations, the water capacity of pools was not a key variable in predicting *D. tinctorius* tadpole presence, corroborating Rojas’ (2014) findings. While a higher water-holding capacity is expected to decrease desiccation risk, pool volume and depth are not always reliable measures for water-holding persistence in phytotelmata (Rudolf and Rödel 2005), making frogs adjust their preference based on other pool traits (see below). The presence of large conspecifics, for instance, may be used as a cue of pool stability and thus influence pool choice by males (Rojas 2014).

An unexpected variable that segregated all three frog species was pool salinity, which tends to increase with height. Some high arboreal pools were particularly deep and a low turnover of stagnant water could explain high salinity levels, where ions (and similarly, salts) concentrate in pools over a long period of time (Sawidis et al. 2011). These pools were mostly occupied by *D. tinctorius* tadpoles, which appear to withstand salinity conditions of up to 1,000 ppm. *Allobates femoralis* tadpoles, in contrast, were only found in low-salt environments. Low salinity is likely a byproduct of the high amount of leaf litter, which appears to buffer salt concentrations (Roache et al., 2006; see Supp. Fig 3). As microbial activity is limited by high salinities, low-salt pools are ideal for the production of detritus (Roache et al. 2006), the main food source of *A. femoralis* tadpoles. *Osteocephalus oophagus*.tadpoles were found in salinity ranges from 48 to 225 ppm (Fig. 5C), and the relatively saline pools (> 700 ppm) that occur within *O. oophagus* vertical ranges were only occupied by *D. tinctorius*. Adult *O. oophagus* deposit their egg clutches in the water as opposed to the other two species, which only use phytotelmata as tadpole-rearing sites. The saltier conditions of pools higher in the canopy may not be suitable for eggs (Christy and Dickman 2002, Albecker and McCoy 2017), which may limit the suitable conditions for oviposition in *O. oophagus*. Poison frogs, in contrast, can escape these limitations because their clutches are terrestrial. Although the small sample size does not allow any stronger interpretation, it appears that both chemical and physical variables are important in shaping ideal pool conditions in *A. femoralis* and *O. oophagus*.

### Pool choice and flexibility in *Dendrobates tinctorius*

Over the two sampling seasons, the bulk of our study focused on the factors that shaped *D. tinctorius* presence and persistence. Despite having sampled over 80 unique pools and found 350 *D. tinctorius* tadpoles (N = 208, 2019; N = 142, 2020), understanding the critical variables that drive *D. tinctorius* pool choice is difficult because of the wide range of physical and chemical properties in which these tadpoles occur. Further, the interactions between physical, chemical, and biological characteristics in phytotelmata are complex and collinear. Based on both our principal component regression and analysis, we found that the probability of detecting *D. tinctorius* tadpoles increases positively with specific physical (height), chemical (KH, salinity, and hardness), and biological (invertebrate diversity) properties. Interestingly, we found that salinity, hardness, and KH also tend to increase with increasing height (Supp. Fig 4). Overall, these chemical components tend to vary in the same direction when moving up the vertical axis, suggesting a positive relationship between these chemical and physical traits. Invertebrate diversity of occupied pools, in contrast, tends to stay relatively constant across heights and might serve as an important food source for predatory *D. tinctorius* tadpoles.

In this study, we found that KH increases with height, and pools with high KH are more likely to have tadpoles in them. KH is a measure of a solution’s buffering capacity or, in other words, a solution’s resistance to pH changes (Yang et al. 2008). KH values in low arboreal and ground-access pools usually ranged from 3 to 6 KH, while average KH in high canopy pools was 15. Interestingly, two of the five lower pools with a KH above 8 had *D. tinctorius* counts of over 10 tadpoles, demonstrating that, when these conditions are available terrestrially, *D. tinctorius* fathers take advantage of them. The apparent preference for high alkalinity environments is interesting, as work studying the formation of fungal granules has established that high-alkalinity conditions inhibit fungal growth (Yang et al. 2008). The potential relationship of KH limiting the growth of fungi in phytotelm conditions is noteworthy as amphibian fungal pathogens such as *Batrachochytrium dendrobatidis* (*Bd*) spread aquatically (Rosenblum et al. 2010), and *Bd* presence (prevalence of ∼5%) has been reported for *D. tinctorius* in our study area (Courtois et al. 2015). Thus, the consistent detection of *D. tinctorius* tadpoles in high KH pools could indicate that fathers are selecting environments less prone to fungal contamination. Although we are unsure of the proximate mechanisms driving the fathers’ choice of particular chemical conditions in phytotelmata, we establish here that KH, hardness, and salinity play an important role in shaping *D. tinctorius* pool choice, and suspect that these chemical conditions may be linked to the long turn-over time of high arboreal pools.

### The stability of ephemeral pools

In 2020, we were able to follow a subset of low arboreal and ground access pools over a month, recording the pH and *D. tinctorius* tadpole presence on a weekly basis. We found that pools made of dead substrate (fallen palm bracts, dead trees) had a higher pH than live substrates (tree holes). The gross average pH of dead phytotelmata across our sample was 5.68, which is almost exactly the value of unpolluted rainwater (pH = 5.65 when saturated with atmospheric CO_2_; (Koshy et al. 1997)). In contrast to most live substrates (average pH = 4.35), dead phytotelmata are usually in canopy gaps, where rain falls directly into the pools. When reported, the pH of most phytotelmata is acidic (Kitching 2001, von May et al. 2009, Poelman et al. 2013, Ramos et al. 2017); but see Lehtinen 2004, which shows that *Pandus* bromeliads were close to neutral pH). However, most studies on phytotelmata are biased towards living plants and trees, and assess chemical/biological variables of pools at a single time point.

Throughout the month, all pool types decreased in pH (Table 2); a similar trend was also found in bamboo phytotelmata in Peru (von May et al. 2009), suggesting a time-dependent process causing pools to become increasingly acidic over time. Remarkably, some pools dried out multiple times during our sampling period, and when refilled by rain were approximately at the same pH as before the drying event (ex. palm bract originally pH 6.98 (Week 1), dried out (Week 2), refilled pH 6.87 (Week 3); live tree hole pH 2.91 (Week 1), dried out (Week 2), pH 3.02 (Week 3)). This indicates that pool substrate may play an important role in establishing pool pH. Three out of the five pools that dried out were dead palm bracts, suggesting that this pool type, despite having favourable chemical conditions when filled, may also be at a higher risk for desiccation and decomposition.

Surprisingly, pools in dead substrates, such as palm bracts and fallen trees, contained more tadpoles than other pool types despite drying out more regularly across our month survey (Table 3). Such pools tend to occur in forest gaps, which makes them more prone to dessication. However, pools in these lit areas may also have more microbial activity and less food limitation (Kitching 2001, Rudolf and Rödel 2005), making them attractive deposition sites for tadpoles. Suitable pools are a limiting resource for frogs and other animals (Donnelly 1989a, b, Fincke 1992, Poelman and Dicke 2007, Ringler et al. 2015) and new pools for *D. tinctorius*, such as those in tree-fall gaps, appear unpredictably and are rapidly occupied despite the high rates of competition and cannibalism (Rojas 2015). This strategy can be particularly beneficial when parents arrive early to new pools, as it allows their offspring to be predators rather than prey. Thus, the competition to be the first to deposit tadpoles might make pools in dead substrates that occasionally dry out additionally attractive.

Interestingly, the size range of tadpoles in dead substrates is much more variable than in low and high arboreal pools (CF, BR, AP pers. observ.), suggesting that the pools remain attractive even when already occupied by larger cannibals. This pattern corroborates the experimental evidence that *D. tinctorius* preferably deposit newly-hatched tadpoles in pools already occupied by conspecifics (Rojas 2014, 2015); possibly, tadpole presence serves as an indicator of pool stability, which might be more valuable to a father’s deposition choice than the risk of having his offspring cannibalized by conspecifics.

### High arboreal pools

While most of our work focused on low arboreal and ground access pools, this study provides one of the first thorough characterisations of high arboreal phytotelmata in the Amazon. Gaucher (2002) unexpectedly found *D. tinctorius* tadpoles in canopy pools up to 25 meters high while studying the treefrog *Trachycephalus hadroceps*. Other poison frogs, such as *D. auratus* have been reported to use arboreal tree holes as well (Summers 1990). We found large numbers of tadpoles in arboreal pools up to 20 m in height, which suggests some benefit of these pools given the presumed high energetic expense that fathers invest in transporting their tadpoles from terrestrial oviposition sites.

One key advantage of high arboreal phytotelmata may be a regular food source provided by *Trachycephalus* treefrogs that specialize in these pools. During this study, all of the suitable high arboreal pools were found by locating nocturnal calls of *T. resinifictrix and T. hadroceps*, indicating that these habitats were potentially used as breeding sites. Although the breeding frequency of these treefrog species is sporadic (Gaucher 2002), successful breeding events result in clutches that consist of hundreds to thousands of eggs and tadpoles, which *D. tinctorius* tadpoles readily consume ((Gaucher 2002), personal obs. AP and BR). As proposed by Gaucher (2002), it could be that *D. tinctorius* fathers cue on *Trachycephalus* calls for locating high arboreal pools, but this warrants further investigation.

Unexpectedly, we also found that dead substrate pools share some characteristics with high arboreal pools, particularly with respect to chemical qualities (a more basic pH), tadpole abundance, and being a limited or hard-to-access resource. As such, both pool types offer benefits that fathers may value: despite having a shorter life, novel pools (such as fallen palm bracts and holes in fallen trees) are worth invading as deposition sites because they are easy to access and have a high probability of having food, and a suitable chemical profile; high arboreal pools, on the other hand, may have sporadic food and are hard to access, but they are stable and less prone to chemical fluctuations. Together, these different pools are both worthy deposition sites, as they provide different stable environments which create a range of possible offspring success.

## Conclusions

When comparing the occurrence of tadpole species in pools, one of the first trends that emerges is the presence/absence on the basis of specific phytotelm characteristics. For example, *A. femoralis* and *O. oophagus* vertical ranges technically overlap, yet tadpoles never co-occur. In species that demonstrate a distinct morphological limitation or vertical preference, it may be that tadpoles occur in pools because that is what is available to their parents. These constraints play a role in the environment tadpoles are exposed to, and should affect their physiology and behavior. But what about when parents are completely unconstrained? *Dendrobates tinctorius* occur across the vertical gradient and occupy pools that range from acidic (pH = 2.96) to neutral pH, with volumes from 19 mL to over 270L and in pools that range from fresh to slightly saline (∼1000 ppm), which hints at a remarkable physiological flexibility that has been overlooked thus far. Therefore, physiological studies comparing phytotelm-breeding tadpoles would be especially interesting to better understand parental decisions. It is also warranted to measure *D. tinctorius* growth in pools with different chemical compositions to see if (despite surviving) these tadpoles are paying a cost for the deposition choices by their fathers.

In sum, natural history studies allow us to grasp species’ flexibility; this is becoming increasingly relevant when we consider the effects of climate change in the Amazon. Forecasted changes in precipitation (Cochrane and Barber 2009, Silvestrini et al. 2011) will have direct effects on animals dependent on plant-based water bodies, and surely the chemical composition of the water bodies themselves. Further studies examining larval anuran responses to challenging environmental conditions (especially KH and pH) are necessary to better understand the potential resistance and adaptability of phytotelm-dependent frogs and how this may shape species resilience in the future.

## Significance statement

This two-year field project is a reflection of over a decade of natural history observation and experiments in the Neotropics. In this study, we expand our knowledge of the ecology of phytotelm-dependent frogs, this time focusing on a comparative overview between larval species and the precise ecological factors that shape the microhabitats in which eggs and larvae are deposited. The breadth of this work will surely engage a wide variety of readers who are interested in ecological dynamics in the tropics.

Here, we synthesize the occurrence and interactions between three different amphibian species, which contributes to understanding the dynamics of each species independently, but this work is also a perspective into the interactions between larvae and parents within the context of an ecological study. Species in this study are specifically plant-dependent as tadpoles, which provides a unique opportunity to thoroughly survey and measure the microhabitats in which they occur. Together, our findings show how biological, physical, and chemical components interact in predicting larval presence in species with parental care, which is a strategy present in ten percent of amphibians. We also provide the first detailed account of the ecology of high arboreal breeding pools. Accessing trees more than 20 meters in height is challenging, and conducting a thorough survey of these environments framed in comparison to microhabitats across the vertical gradient is a testament to the scope of this work. While our study is based on amphibians, it more broadly focuses on the ecology that shapes larval deposition sites and the wide array of species’ flexibility we observe in the Neotropics.

Hopefully, a wide variety of researchers will be excited to learn more about the diversity of microhabitats in the Amazon and a cross-species comparison of the amphibians that depend on them.

## Data availability statement

All data will be archived in the data repository of the University of Jyväskylä.

## Acknowledgements

We are grateful to the staff of Nouragues Ecological Research Station (managed by CNRS), which benefits from “Investissement d’Avenir” grants managed by Agence Nationale de la Recherche (AnaEE France ANR-11-INBS-0001; Labex CEBA ANR-10-LABX-25-01), for logistic support in the field and for providing the meteorological data. This work is part of a partnership between BR, AP, and the Nouragues Nature Reserve aimed at improving and spreading the knowledge about amphibians. We thank the staff of the Nouragues Nature Reserve for their commitment to preserving our natural world; Walter Hödl for his ongoing mentorship and inspiration of this work and collaboration; and Lauren A. O’Connell for her generous support of AP, SJSR, and MTF. A huge *grazie mille* to Matteo Vecchi for being patient in teaching CF how to truly attack a PCA with all of the statistical might humanly possible. The authors highly value equity, diversity and inclusion in science. We cherish the international and diverse nature of our team, which includes researchers from (7) different countries, backgrounds and career stages, as it significantly contributed to the fulfilment and quality of the present study.

## Ethics statement

The study was approved by the scientific committee of the Nouragues Ecological Research Station and covered under a partnership agreement between BR, AP and the Nouragues Nature Reserve (No 01-2019). We strictly adhered to the current French and European Union law, and followed the Association for the Study of Animal Behaviour’s (ASAB) Guidelines for the use of live animals in teaching and research (ASAB, 2017).

## Author contributions

**Chloe Fouilloux:** Writing – original draft (lead); Investigation (equal); Data curation (equal); Formal analysis (lead); **Shirley Jennifer Serrano-Rojas:** Investigation (equal); Data curation (equal); Formal analysis (supporting); Writing - review and editing (supporting); **Juan David Carvajal-Castro:** Investigation (equal); Writing - review and editing (supporting); **Marie- Therese Fischer:** Investigation (supporting); Writing - review and editing (supporting); **Janne Valkonen:** Methodology (supporting); Investigation (supporting); Writing - review and editing (supporting); **Philippe Gaucher:** Investigation (supporting); **Andrius Pašukonis:** Conceptualization (equal); Methodology (equal); Investigation (equal); Writing – review and editing (equal); Supervision (equal); Funding acquisition (equal); **Bibiana Rojas:** Conceptualization (equal); Methodology (equal); Investigation (supporting); Writing – review and editing (equal); Supervision (equal); Funding acquisition (equal).

## Speculations and alternative viewpoints

*Dendrobates tinctorius* males typically father egg clutches of 2-5 tadpoles pers clutch and breed year-round (Rojas and Pašukonis 2019). As a result of the presumed high energetic expense from carrying each tadpole from each clutch singly, we hypothesize that tadpole transported later may be subject to bet-hedging by fathers.

**Fig xx.**
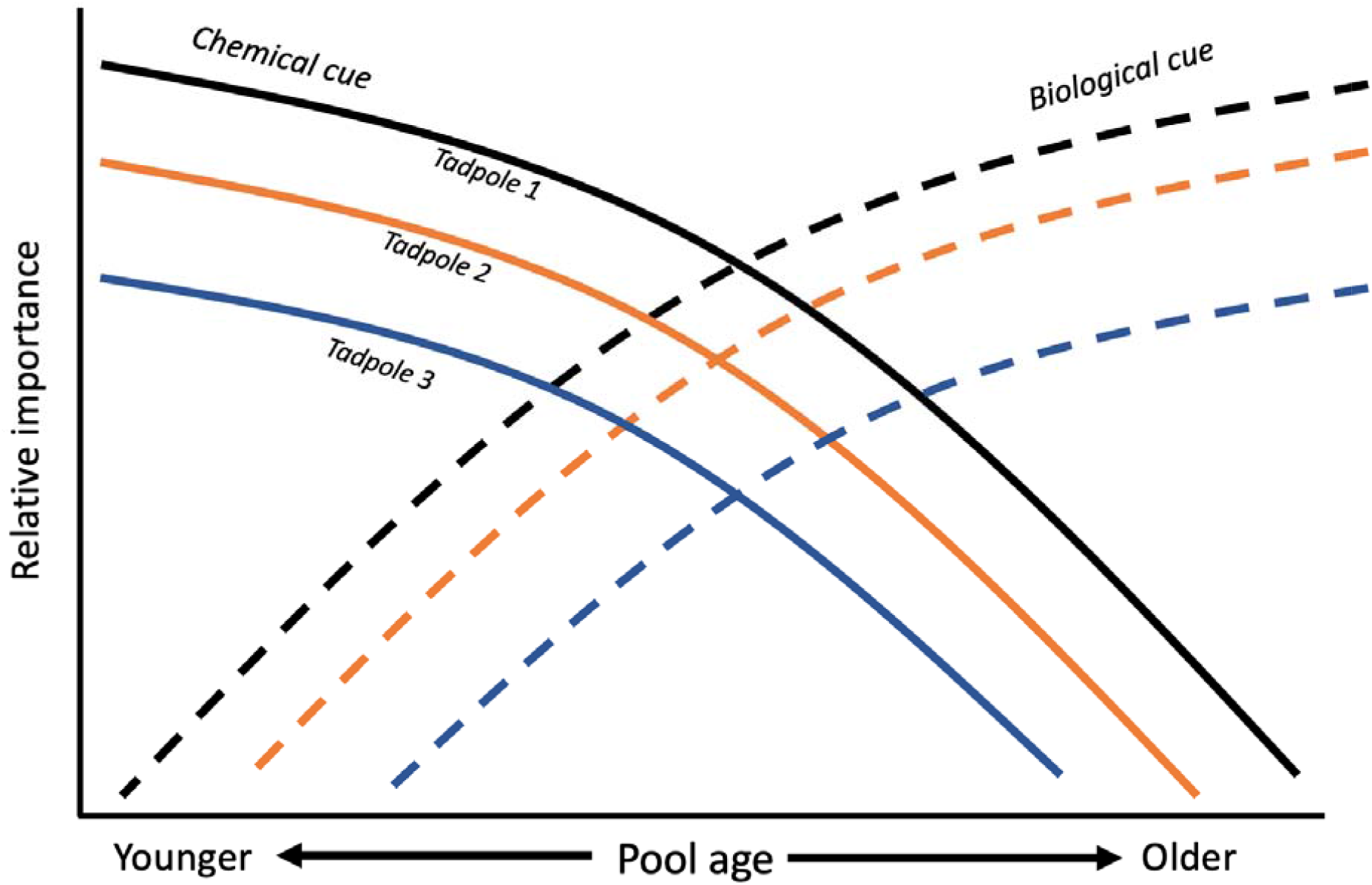
Hypothetical succession of relative cue importance in phytotelmata across time.

Combined with the important chemical aspects of pools shown from 2019 data, it seems that fathers can cue on either chemical (KH, salinity, hardness) or biological components (the presence of conspecifics) as information about pool stability. We speculate that the importance of chemical and physical cues changes with respect to pool age. For example, in new uninvaded pools, chemical cues of the pool may be more important (left-side of figure), while in older, established pools that are more densely occupied, density serves as a main cue for transporting fathers (right-side of figure). Finally, the value of these cues may vary with the amount of offspring fathers are caring for. Hypothetically, fathers who must transport more offspring are less discerning of where they transport latter tadpoles because they can afford to make less “ideal” deposition decisions because of their large reproductive output.

## References

Akaike, H. 1974. A new look at the statistical model identification. - IEEE Trans. Automat. Contr. 19: 716–723.

Albecker, M. A. and McCoy, M. W. 2017. Adaptive responses to salinity stress across multiple life stages in anuran amphibians. -Front. Zool. 14: 40.

Almendáriz, A. et al. 2000. Two new species of hylid frogs, genus Osteocephalus, from Amazonian Ecuador. -Amphib-Reptil. 21: 327–340.

Björklund, M. 2019. Be careful with your principal components. - Evolution 73: 2151–2158.

Brown, C. R. and Brown, M. B. 1991. Selection of high-quality host nests by parasitic cliff swallows. - Anim. Behav. 41: 457–465.

Brown, G. P. and Shine, R. 2005. Nesting snakes (Tropidonophis mairii, Colubridae) selectively oviposit in sites that provide evidence of previous successful hatching. - Can. J. Zool. 83: 1134–1137.

Brown, J. L. et al. 2008a. Phytotelm size in relation to parental care and mating strategies in two species of Peruvian poison frogs. - Behaviour 145: 1139–1165.

Brown, J. L. et al. 2008b. Divergence in parental care, habitat selection and larval life history between two species of Peruvian poison frogs: an experimental analysis. - J. Evol. Biol. 21: 1534–1543.

Brown, J. L. et al. 2010. A key ecological trait drove the evolution of biparental care and monogamy in an amphibian. - Am. Nat. 175: 436–446.

Bult, A. and Lynch, C. B. 1997. Nesting and fitness: lifetime reproductive success in house mice bidirectionally selected for thermoregulatory nest-building behavior. - Behav. Genet. 27: 231–240.

Christy, M. and Dickman, C. 2002. Effects of salinity on tadpoles of the green and golden bell frog (Litoria aurea). - Amphib-reptil. 23: 1–11.

Cochrane, M. A. and Barber, C. P. 2009. Climate change, human land use and future fires in the Amazon. - Glob. Chang. Biol. 15: 601–612.

Core, R. 2015. Team. - R: a language and environment for statistical computing 3: 2. Courtois, E. A. et al. 2015. Widespread occurrence of bd in French Guiana, South America. -PLoS One 10: e0125128.

Crump, M. L. 1974. Reproductive strategies in a tropical anuran community. - University of Kansas Museum of Natural History Miscellaneous Publications 61: 1–68.

Donnelly, M. A. 1989a. Demographic effects of reproductive resource supplementation in a territorial frog, Dendrobates pumilio. - Ecol. Monogr. 59: 207–221.

Donnelly, M. A. 1989b. Effects of reproductive resource supplementation on space-use patterns in Dendrobates pumilio. - Oecologia 81: 212–218.

Ebensperger, L. A. et al. 2014. Mean ecological conditions modulate the effects of group living and communal rearing on offspring production and survival. - Behav. Ecol. 25: 862–870.

Fincke, O. M. 1992. Interspecific Competition for Tree Holes: Consequences for Mating Systems and Coexistence in Neotropical Damselflies. - Am. Nat. 139: 80–101.

Gaucher, P. 2002. Premières données sur Phrynohyas hadroceps, Rainette arboricole du plateau des Guyanes (Amphibia:Anura:Hylidae) (Révision taxonomique, éco-éthologie de la reproduction).

Gomez-Mestre, I. et al. 2012. Phylogenetic analyses reveal unexpected patterns in the evolution of reproductive modes in frogs. - Evolution 66: 3687–3700.

Gray, H. M. et al. 2009. Kin discrimination in cannibalistic tadpoles of the Green Poison Frog, Dendrobates auratus (Anura, Dendrobatidae). - Phyllomedusa 8: 41–50.

Hartig, F. 2020. DHARMa: Residual Diagnostics for Hierarchical (Multi-Level/Mixed) Regression Models (2017). - R package version 0. 1 in press.

Jungfer, K.-H. and Weygoldt, P. 1999. Biparental care in the tadpole-feeding Amazonian treefrog Osteocephalus oophagus. - Amphib-reptil. 20: 235–249.

Kitching, R. L. 2001. Food webs in phytotelmata: “bottom-up” and “top-down” explanations for community structure. - Annu. Rev. Entomol. 46: 729–760.

Koshy, K. et al. 1997. Wet deposition chemistry studies at Suva, Fiji, a remote tropical island site in the south Pacific. - Environ. Geochem. Health 19: 0–0.

Lehtinen, R. M. 2004. Tests for competition, cannibalism, and priority effects in two phytotelm-dwelling tadpoles from Madagascar. - Herpetologica 60: 1–13.

Lin, Y.-S. et al. 2008. Time- and Context-Dependent Oviposition Site Selection of a Phytotelm-Breeding Frog in Relation to Habitat Characteristics and Conspecific Cues. - Herpetologica 64: 413–421.

Magnusson, W. E. and Hero, J.-M. 1991. Predation and the evolution of complex oviposition behaviour in Amazon rainforest frogs. - Oecologia 86: 310–318.

Magnusson, A. et al. 2020. Package “glmmTMB”: Generalized Linear Mixed Models using Template Model Builder. The Comprehensive R Archive Network. in press.

Marsh, D. M. and Borrell, B. J. 2001. Flexible oviposition strategies in túngara frogs and their implications for tadpole spatial distributions. - Oikos 93: 101–109.

McKeon, C. S. and Summers, K. 2013. Predator driven reproductive behavior in a tropical frog. - Evol. Ecol. 27: 725–737.

Mikheev, V. N. et al. 2001. Spatial distribution and hatching of overwintered eggs of a fish ectoparasite, Argulus coregoni (Crustacea: Branchiura). - Dis. Aquat. Organ. 46: 123– 128.

Mokany, A. and Shine, R. 2003. Oviposition site selection by mosquitoes is affected by cues from conspecific larvae and anuran tadpoles. - Austral Ecol. 28: 33–37.

Narins, P. M. et al. 2003. Bimodal signal requisite for agonistic behavior in a dart-poison frog, Epipedobates femoralis. - Proc. Natl. Acad. Sci. U. S. A. 100: 577–580.

Nussbaum, R. A. 1987. Parental care and EGG size in salamanders: An examination of the safe harbor hypothesis. - Res. Popul. Ecol. 29: 27–44.

Ottesen, O. H. and Bolla, S. 1998. Combined effects of temperature and salinity on development and survival of Atlantic halibut larvae. - Aquac. Int. 6: 103–120.

Pašukonis, A. et al. 2017. Induced parental care in a poison frog: a tadpole cross-fostering experiment. - J. Exp. Biol. 220: 3949–3954.

Pašukonis, A. et al. 2019. How far do tadpoles travel in the rainforest? Parent-assisted dispersal in poison frogs. - Evol. Ecol. 33: 613–623.

Pettitt, B. A. et al. 2018. Predictors and benefits of microhabitat selection for offspring deposition in golden rocket frogs. - Biotropica 50: 919–928.

Poelman, E. H. and Dicke, M. 2007. Offering offspring as food to cannibals: oviposition strategies of Amazonian poison frogs (Dendrobates ventrimaculatus). - Evol. Ecol. 21: 215–227.

Poelman, E. H. et al. 2013. Amazon poison frogs (Ranitomeya amazonica) use different phytotelm characteristics to determine their suitability for egg and tadpole deposition. - Evol. Ecol. 27: 661–674.

Ramos, G. J. P. et al. 2017. Algae in phytotelmata from Caatinga: first record of the genus Rhopalosolen Fott (Chlorophyta) for Brazil. - Check List 13: 403–410.

Ringler, M. et al. 2009. Site fidelity and patterns of short- and long-term movement in the brilliant-thighed poison frog Allobates femoralis (Aromobatidae). - Behav. Ecol.Sociobiol. in press.

Ringler, E. et al. 2013. Tadpole transport logistics in a Neotropical poison frog: indications for strategic planning and adaptive plasticity in anuran parental care. - Front. Zool. 10: 67.

Ringler, M. et al. 2015. Populations, pools, and peccaries: simulating the impact of ecosystem engineers on rainforest frogs. - Behav. Ecol. 26: 340–349.

Ringler, E. et al. 2018. Hierarchical decision-making balances current and future reproductive success. - Mol. Ecol. 27: 2289–2301.

Roache, M. C. et al. 2006. Effects of salinity on the decay of the freshwater macrophyte, Triglochin procerum. - Aquat. Bot. 84: 45–52.

Roithmair, M. E. 1992. Territoriality and male mating success in the dart-poison frog, Epipedobates femoralis (dendrobatidae, Anura). - Ethology 92: 331–343.

Rojas, B. 2014. Strange parental decisions: fathers of the dyeing poison frog deposit their tadpoles in pools occupied by large cannibals. - Behav. Ecol. Sociobiol. in press.

Rojas, B. 2015. Mind the gap: treefalls as drivers of parental trade-offs. - Ecol. Evol. 5: 4028– 4036.

Rojas, B. and Pašukonis, A. 2019. From habitat use to social behavior: natural history of a voiceless poison frog, Dendrobates tinctorius. - PeerJ 7: e7648.

Rosenblum, E. B. et al. 2010. The deadly chytrid fungus: a story of an emerging pathogen. - PLoS Pathog. 6: e1000550.

Ruano-Fajardo, G. et al. 2014. Bromeliad selection by two salamander species in a harsh environment. - PLoS One 9: e98474.

Rudolf, V. H. W. and Rödel, M.-O. 2005. Oviposition site selection in a complex and variable environment: the role of habitat quality and conspecific cues. - Oecologia 142: 316–325.

Sawidis, T. et al. 2011. Trees as bioindicator of heavy metal pollution in three European cities. - Environ. Pollut. 159: 3560–3570.

Schulte, L. M. et al. 2011. The smell of success: choice of larval rearing sites by means of chemical cues in a Peruvian poison frog. - Anim. Behav. 81: 1147–1154.

Schulte, L. M. et al. 2020. Developments in Amphibian Parental Care Research: History, Present Advances, and Future Perspectives. - Herpetological Monographs. 34: 71–97.

Sih, A. and Moore, R. D. 1993. Delayed hatching of salamander eggs in response to enhanced larval predation risk. - Am. Nat. 142: 947–960.

Silvestrini, R. A. et al. 2011. Simulating fire regimes in the Amazon in response to climate change and deforestation. - Ecol. Appl. 21: 1573–1590.

Summers, K. 1990. Paternal care and the cost of polygyny in the green dart-poison frog. - Behav. Ecol. Sociobiol. 27: 307–313.

Summers, K. and McKeon, C. S. 2004. The evolutionary ecology of phytotelmata use in Neotropical poison frogs. - Miscellaneous Publications, Museum of Zoology, University of Michigan 193: 55–73.

Summers, K. and Tumulty, J. 2014. Chapter 11 - Parental Care, Sexual Selection, and Mating Systems in Neotropical Poison Frogs. - In: Macedo, R. H.and Machado, G. (eds), Sexual Selection. Academic Press, pp. 289–320.

Svendsen, G. E. 1976. Structure and Location of Burrows of Yellow-Bellied Marmot. - Southwest. Nat. 20: 487–494.

Touchon, J. C. and Worley, J. L. 2015. Oviposition site choice under conflicting risks demonstrates that aquatic predators drive terrestrial egg-laying. - Proc. Biol. Sci. 282: 20150376.

Vági, B. et al. 2019. Parental care and the evolution of terrestriality in frogs. - Proc. Biol. Sci. 286: 20182737.

von May, R. et al. 2009. Breeding-site selection by the poison frog Ranitomeya biolat in Amazonian bamboo forests: an experimental approach. - Can. J. Zool. 87: 453–464.

Warkentin, K. M. 2011. Environmentally cued hatching across taxa: embryos respond to risk and opportunity. - Integr. Comp. Biol. 51: 14–25.

Wells, K. D. 2007. The Ecology and Behavior of Amphibians. - University of Chicago Press.

Weygoldt, P. 1980. Complex brood care and reproductive behaviour in captive poison-arrow frogs, Dendrobates pumilio O. Schmidt. - Behav. Ecol. Sociobiol. 7: 329–332.

Williams, B. K. et al. 2008. Leaf litter input mediates tadpole performance across forest canopy treatments. - Oecologia 155: 377–384.

Yang, S. F. et al. 2008. Formation and characterisation of fungal and bacterial granules under different feeding alkalinity and pH conditions. - Process Biochem. 43: 8–14.

Zhao, Q.-S. et al. 2016. Nest site choice: a potential pathway linking personality and reproductive success. - Anim. Behav. 118: 97–103.

