## Supplementary Table 1 for "Pool choice in a vertical landscape: tadpole rearing site flexibility in phytotelm-breeding frogs"

**Supp. Table 1.** Definition of variables (traits) considered in the principal component analysis

| Variable | Category | Description |
| --- | --- | --- |
| Salinity | Chemical | Quantification of salt in solution. Range = 0-10000 ppm. |
| KH (Alkalinity) | Chemical | Quantification of pool bicarbonate/carbonate in solution. Range = 0-25 dKH |
| NO^-^_3_ | Chemical | Quantification of nitrate in solution. Range = 0-160 ppm. |
| Hardness | Chemical | Quantification of ions in solution (e.g. calcium). Range = 0-425 ppm. |
| Height | Physical | Vertical height from the ground to the pool entrance. Measured in cm. |
| Water capacity | Physical | Water holding capacity of the pool. Estimated from pool width, length and depth using a semi-ellipsoid formula. |
| Surface area to depth ratio | Physical | Surface area to depth ratio. Surface area calculated from semi-ellipsoid formula. |
| Leaf litter volume | Physical | The measure of leaf litter volume in each pool. |
| Amphibian diversity | Biological | The sum of all species observed using each pool including adults, calling, dead tadpoles and opportunistic observations after the sampling. |
| Invertebrate density | Biological | Sum of all invertebrate densities (counts divided by sampling volume) |
| Invertebrate diversity | Biological | Number of distinct invertebrate categories observed in each pool (between 0 and 12) |
| Predator count | Biological | Number of Odonata larvae in each pool. |
| Average predator size | Biological | The average size of Odonata larvae in each pool. Size is calculated by dividing (pred_size_sum)/(pred_count) |
| Total other | Biological | Sum of *O. oophagus* and *A. femoralis* tadpoles co-occurring in the pool. |
