## Supplementary Table 2 for "Pool choice in a vertical landscape: tadpole rearing site flexibility in phytotelm-breeding frogs"

**Supp. Table 2.** PCA model analysis rank using AIC. All models were coded with a negative binomial family in a GLM framework. Two models fell within 2 AIC of each other; the interaction in the second model was not significant, so we chose the simplest model of the two (Rank 1).

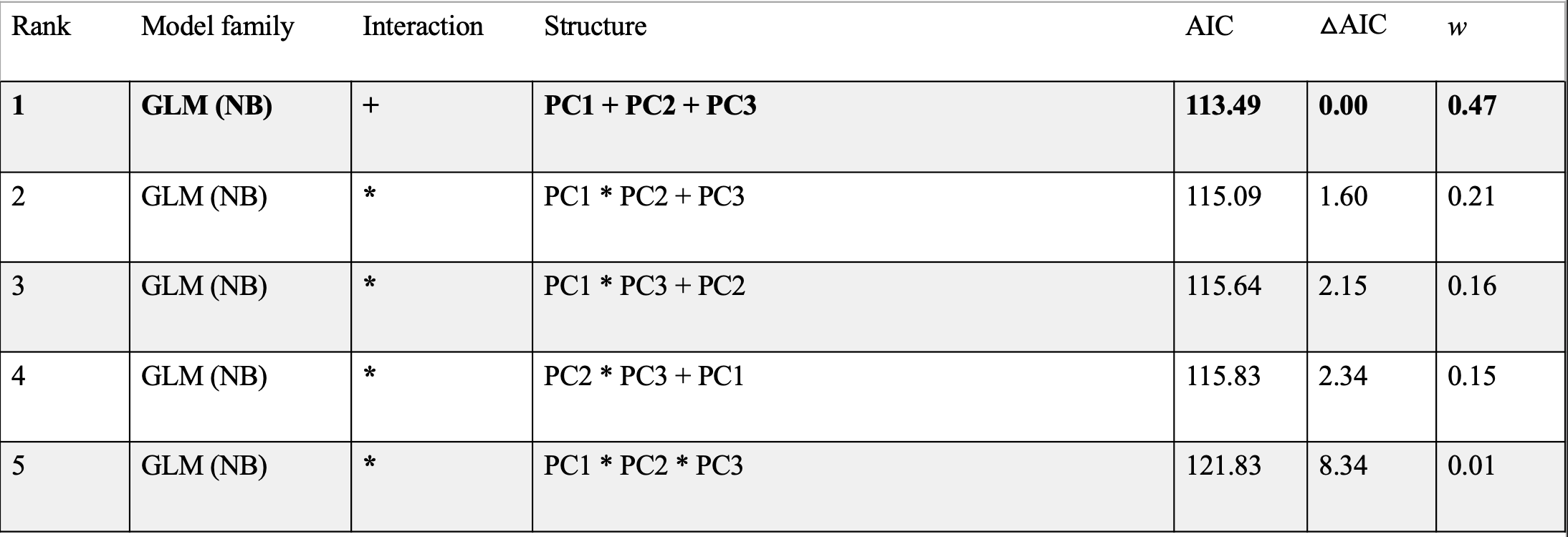
