## Supplementary Table 3 for "Pool choice in a vertical landscape: tadpole rearing site flexibility in phytotelm-breeding frogs"

**Supp. Table 3.** **drop1 model selection for the predictors of *pH* measure after repeated pool observations (2020 data)**. Bolded components of each row indicate dropped part of each model iteration. Pool type was a binomial categorical variable (dead/alive), Week indicates week of sampling (numerical variable: 1-4), water capacity was a continuous variables (depth*length*width) of each pool, and Dt_Tadpole_Num was a continuous whole number of *Dendrobates tinctorius* tadpole counts. Interaction term indicated the interaction between covariates. Model family was coded as “Gaussian” for all pH models. Models are ranked in decreasing AIC order. Random effect of pool ID was included in all models.

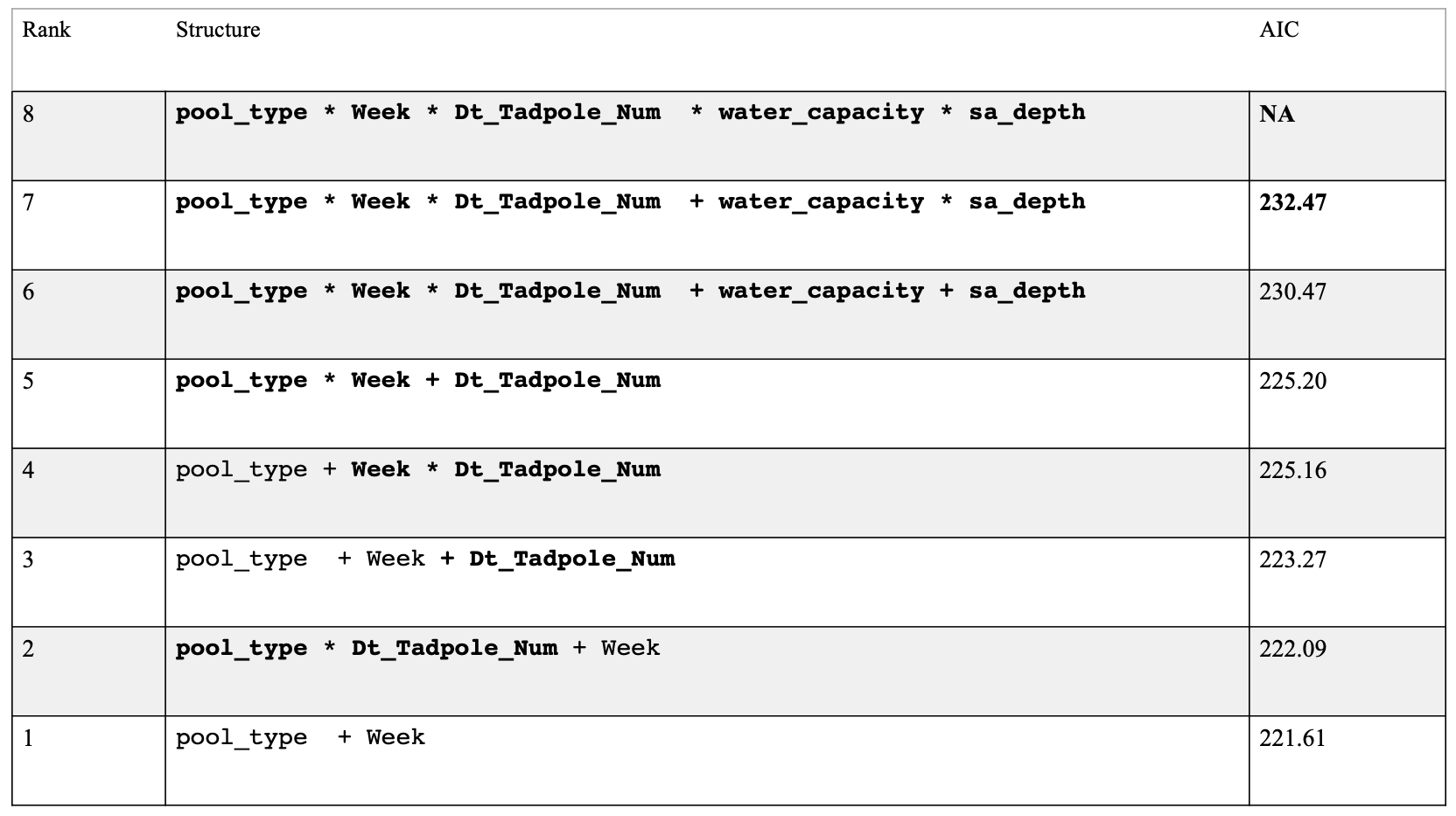
