## Supplementary Table 4 for "Pool choice in a vertical landscape: tadpole rearing site flexibility in phytotelm-breeding frogs"

Supp. Table 4. **drop1 model selection for the predictors of *Dendrobates tinctorius* tadpole numbers after repeated pool observations (2020 data)**. Bolded components of each row indicate dropped part of each model iteration. Pool type was a binomial categorical variable (dead/alive), Week indicates week of sampling (numerical variable: 1-4), and pH was a continuous variable, water capacity is pool volume based on semi-ellipsoid equation, and sa_depth is the surface area to depth ratio of each pool. Interaction term indicated the interaction between covariates. Models are ranked in decreasing AIC value. Random effect of pool ID was included in all models. Models were fit with a quadratic (nbinom2) negative binomial family.

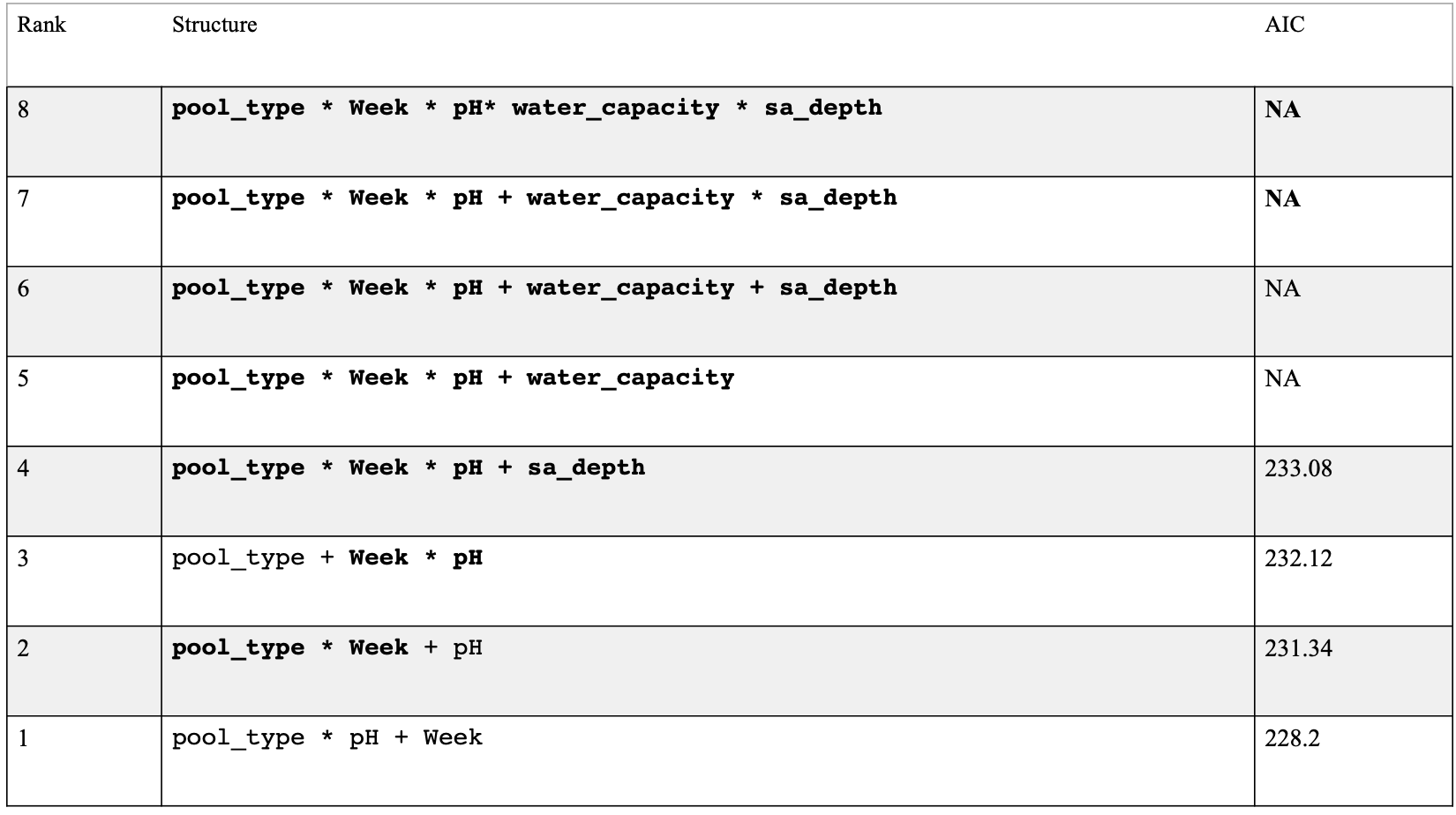
