## Supplementary Figure 1 for "Pool choice in a vertical landscape: tadpole rearing site flexibility in phytotelm-breeding frogs"

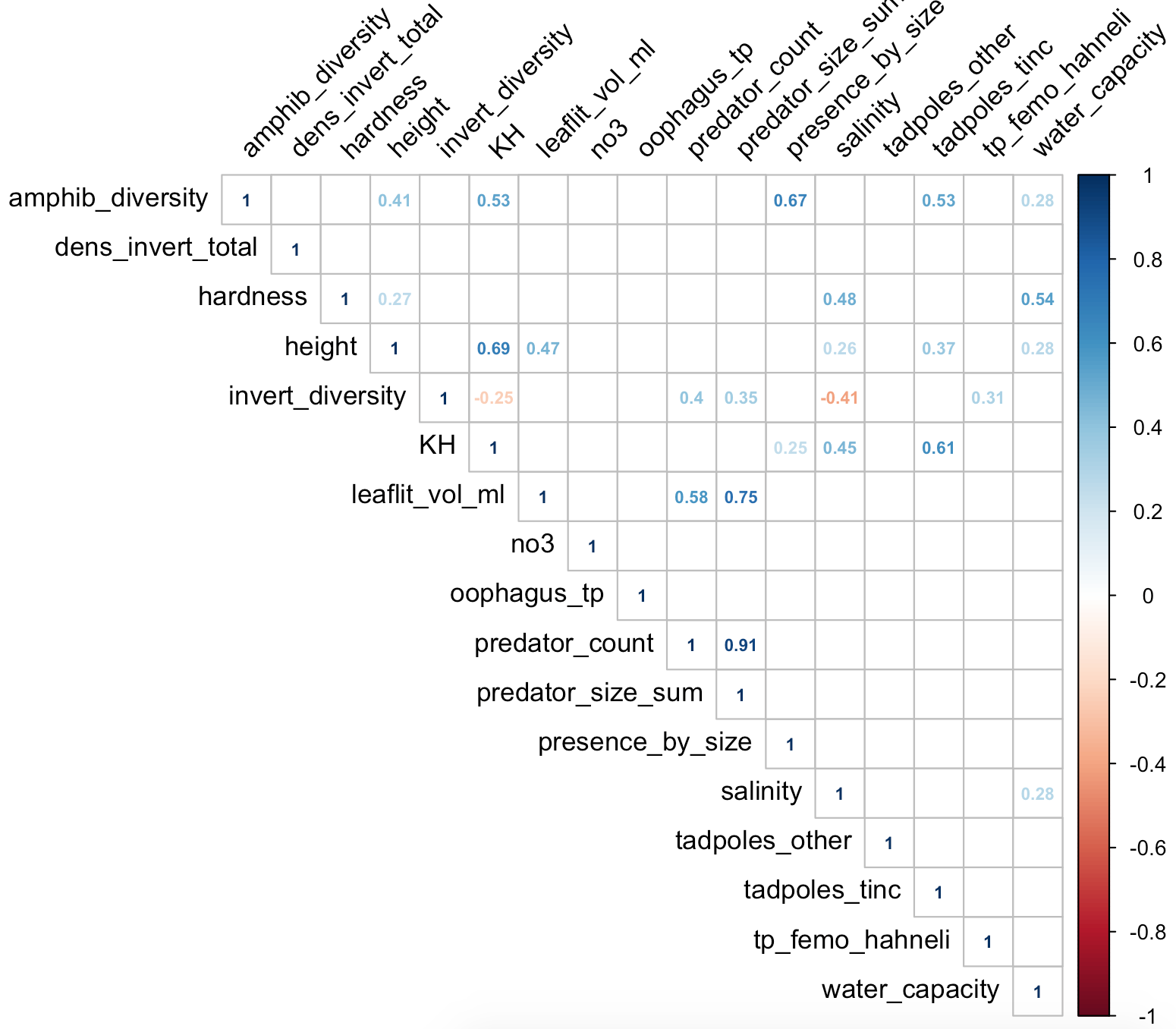

**Supplementary Fig 1. High correlation between numeric variables in frogpool data set (2019).** Only significant correlations (*p* > 0.05) from a Pearsons’s correlation test are visualized. Cooler colors represent positive correlations and warmer colors represent negative correlations. Variables are ordered alphabetically.
