## Supplementary Figure 2 for "Pool choice in a vertical landscape: tadpole rearing site flexibility in phytotelm-breeding frogs"

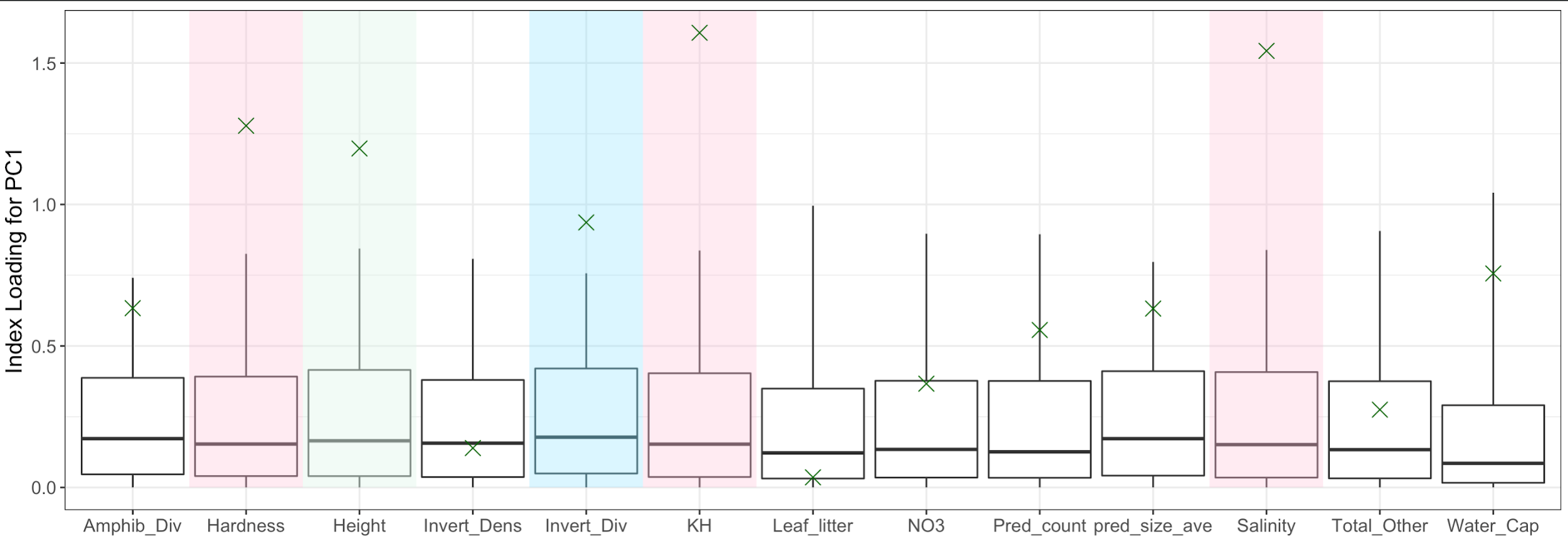

**Supp. Figure 2. Results from the PCA correlation procedure. Boxplots are generated from random data, where whiskers range 95% confidence intervals**; green “X”s are observed PCA index loading values. Variables where the observed PCA index loading are significantly different from the random confidence interval are highlighted. Pink highlight represents chemical variables, green highlight represents physical variables, and blue highlight represent biological variables. P values for each significant variable are shown in Table xx.
