## Supplementary Figure 3 for "Pool choice in a vertical landscape: tadpole rearing site flexibility in phytotelm-breeding frogs"

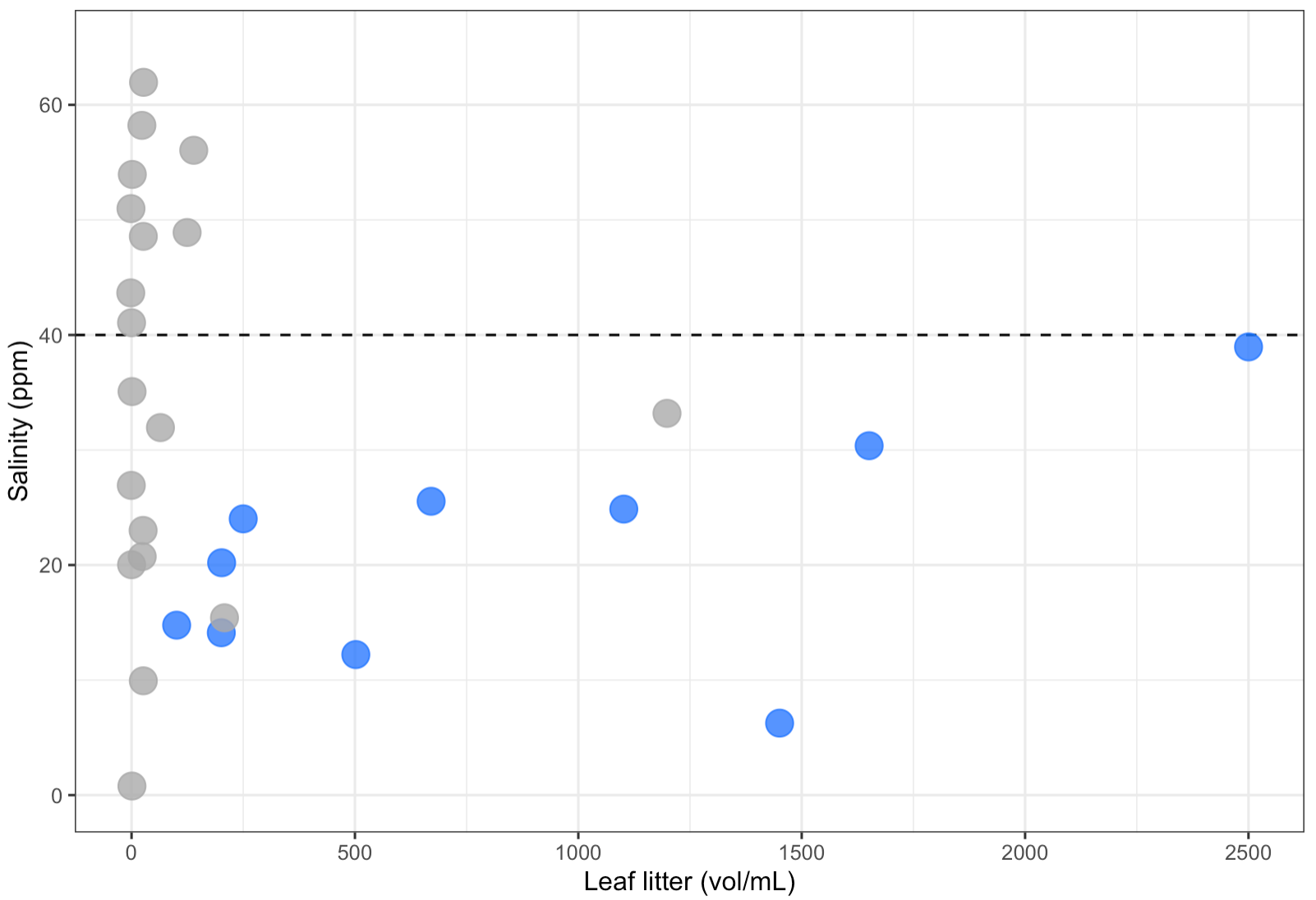

**Supp. Fig 3. Relationship between leaf litter volume and salinity.** Dashed line is at 40 ppm which is the limit where we detected *A. femoralis* tadpoles. Below this level, it appears that leaf litter and salinity have a slightly positive relationship, though interpretation is limited by sample size (blue points, N_Femoralis_= 10). Data is subsetted for ground access pools, and salinity upper bound was limited to 70 ppm to emphasize potential leaf litter effect.
