## Supplementary Figure 4 for "Pool choice in a vertical landscape: tadpole rearing site flexibility in phytotelm-breeding frogs"

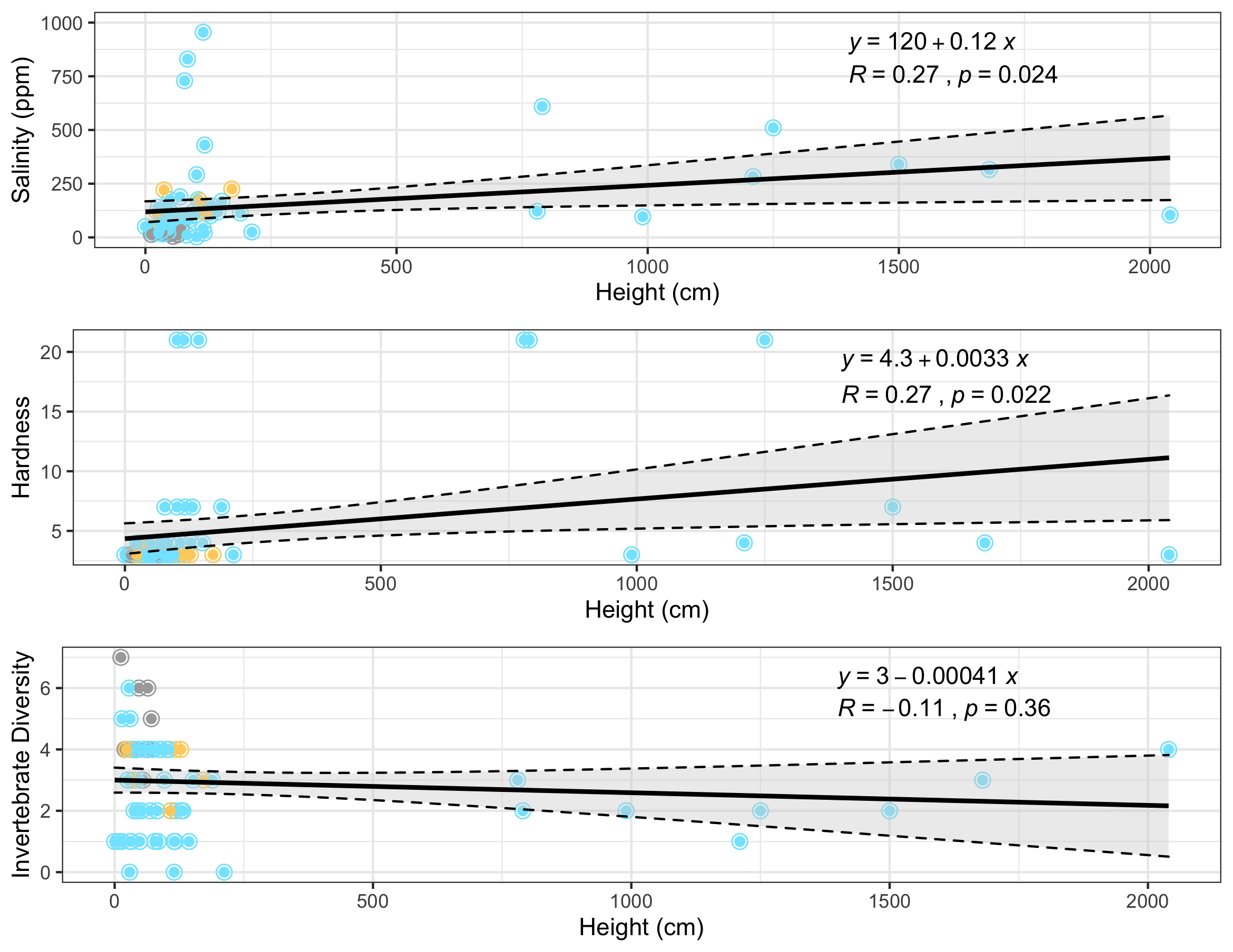

**Supp Fig 4. Relationship between height and salinity, invertebrate diversity, and hardness.** There is a positive relationship between salinity and hardness with height. We see that invertebrate diversity does not meaningfully change with height. Dashed line represents 95% CI. GLM line fitted with a y~x formula.
